# Bound for the nucleus: defining the molecular principles of cargo selection by importin 9

**DOI:** 10.64898/2026.05.12.724649

**Authors:** Amanda J. Keplinger, Tyler C. Cropley, Juliana Ortiz-Pacheco, Prithi A. Srinivasan, Beatrix M. Ueberheide, Alexander J. Ruthenburg

## Abstract

Active nuclear transport of proteins enables essential nuclear processes, such as genome packaging, transcription, splicing, ribosome biogenesis, and DNA repair. To facilitate proper sorting between cytoplasmic and nuclear compartments, the importin class of chaperone proteins transports hundreds of proteins through hydrophobic nuclear pores. While some importins recognize cargos via classical nuclear localization signals (cNLSs), there is not a comprehensive understanding of binding specificity determinants for those that appear to recognize other cargo features. Here we subject one such importin, IPO9, which is known to import H2A-H2B, TFIIB, actin, and the proteasome into the nucleus without cNLS binding, to a detailed analysis of its full set of cargos and the structural elements of IPO9 that confer their recognition. Through cytoplasmic immunoprecipitation followed by mass spectrometry (IP-MS), we stringently and reproducibly identify 79 *bona fide* IPO9-bound cargos, including 20 previously validated cargos. With this comprehensive cargo list, we find that IPO9 does not appear to use cNLSs, nor any other linear peptide motif, to identify and bind cargos. Unbiased oxidative footprinting of extracted IPO9•cargo complexes reveals that both the inner cavity of IPO9 and the unstructured loops protruding from the main body of the importin are protected by bound cargo, indicative of cargo competition for limiting IPO9 capacity. Guided by these data and evolutionary and phosphoproteomics insights, we employ a systematic IP-MS approach with loop perturbations to define how each element contributes to selective binding of subsets of the whole cargo cohort. These data suggest the H8 and H18-19 loops both specifically mediate IPO9 cargo-recognition by favorable enthalpic contacts. Additionally, these loops, as well as H7, appear to preclude binding of a secondary set of potential cargo to IPO9 which may normally be repelled by them or outcompeted by cognate cargo binding for which the loops provide attractive contacts. We define the continuum of cargo-release factor RanGTP sensitivity for our full cargo set, noting orders of magnitude range of sensitivity that argues that additional release factors may be necessary for efficient unloading in the nucleus. These experiments provide structure-function insight into importin binding specificity dictated by structural elements that recruit and/or restrict protein-protein interactions. Our approach of targeted mutations in cellular contexts coupled to quantitative proteomics affords a thorough biochemical dissection of the discrimination principles undergirding this unique molecular recognition problem of numerous-yet-specific binding events between importins and their many distinct cargos.

## Introduction

Cellular localization dictates the ability of proteins to carry out their function (1). The nuclear membrane and hydrophobic nuclear pore channels act as a barrier to delineate protein localization and function, limiting contact of nascent proteins with chromatin. Aberrant nuclear-cytoplasmic sorting of proteins leads to neurodegeneration, cancer, and developmental defects (2–5). Generally, charged proteins or proteins larger than 40 kilodaltons in molecular weight must utilize specialized transport chaperone proteins, known as karyopherins, to traverse the hydrophobic nuclear pores (6–8). In mammals, there are ten karyopherin import proteins: the well-characterized importin β (KPNB1); the transportins TNPO1, TNPO2, TNPO-SR; and the importins IPO4, IPO5, IPO7, IPO8, IPO9, IPO11 (6). Importins are structurally similar, being composed of 18-24 antiparallel alpha helix pairs, HEAT repeats, that form a right-handed solenoid, with a hydrophobic outer face that facilitates movement through nuclear pores (6, 9). Active transport of protein cargos bound by importins relies on a nuclear-to-cytoplasmic gradient of nucleotide-bound Ran, where nuclear RanGTP binding to the importin is thought to induce release of cargo (10, 11). A nuclear-localized chromatin-bound guanine nucleotide exchange factor (RanGEF) reloads GTP onto Ran (12), while a cytoplasmic-side nuclear pore-localized GTPase activating protein (RanGAP) hydrolyzes GTP to GDP while bound to Ran, lowering the affinity of Ran for importins and inducing cytoplasmic dissociation of the importin•Ran complex (13). The propensity of the importin proteins to mediate specific binding, import, and release of hundreds of proteins into the nucleus with fidelity requires a myriad of remarkably complex and dynamic cellular equilibria.

Binding of importins to their numerous cargos remains a relatively unique phenomenon of protein molecular recognition: specific binding to potentially hundreds of proteins is a delicate kinetic balancing act with profound cellular consequences. Despite the structural similarities among the class of importins, importins appear to exhibit binding profiles distinct from one another, occupying cargo niches (14, 15). IMB, IPO7, and IPO8, in concert with smaller importin alpha (IMA) proteins, recognize classical nuclear localization signals (cNLSs) which are characterized by short lysine-rich mono- or bi-partite amino acid motifs (9). Transportins TNPO1 and TNPO2 generally bind to proline and lysine rich PY-NLSs (9, 16). For IPO4, IPO9, IPO11, bioinformatic analyses have been unable to identify cNLSs or other linear mono- or bi-partite motifs present in identified cargos, and several individual structural studies suggest more complex three-dimensional motifs may dictate cargo binding (17–19). Thus, the general principles of cargo recognition and extent to which linear signals mediate binding by this class of importins remain unclear.

Importin 9 (IPO9) is composed of twenty anti-parallel alpha helix HEAT repeat pairs and four major loops extending from this scaffold toward its hollow interior – the H7, H8, H18-19 and H19 loops (20–22). Perturbed expression of IPO9 is implicated in a variety of cancers, Alzheimer’s, and other neurodegenerative diseases (23–29). In mice, homozygous deletion of IPO9 disrupts early development through neuronal, cardiac and patterning morphological defects that culminate in embryonic lethality at day 7 (30), further arguing for its critical role in nuclear proteome homeostasis. Yet Importin 9 is one of the least-studied importins, with identification and dissection of binding modes only occurring for a few cargos (18, 21, 22, 31). Most studies of specific cargo originated in the yeast homolog of IPO9, Kap114 (20, 31–33), including the finding that the central HEAT repeat core is critical for binding and import of the general transcription factor TFIIB (21). A non-conserved C-terminal post-translational modification (PTM) has been shown to be important for Kap114 for binding to Tata Box Binding Protein (TBP) (34). Broadly, IPO9 does not seem to use cNLSs to bind cargos, as the scattered individual cargo interaction studies have shown alternative means of cargo identification and binding (17, 18, 21). We have recently shown that IPO9 binds directly to actin *in vitro*, independent of the cNLS-containing actin binding protein cofilin (35). Binding of the 20S proteasome to IPO9 is mediated by adaptor protein AKIRIN2, which surprisingly contacts the outer surface of the HEAT repeats and does not use a cNLS (17, 22). Biochemical and structural work investigating H2A-H2B histone dimer binding and release by Importin 9 (IPO9) has demonstrated that rather than using the potential cNLS motifs in the H2A or H2B histone N-terminal tails, IPO9 utilizes the unstructured basic H18-19 loop to bind an acidic patch of the dimer (18). Additionally, the HEAT 19 (H19) loop of IPO9 has been shown to be critical for binding Nap1, a critical histone chaperone (36). Though well characterized in these few examples, it remains unclear how widespread unstructured loops are utilized for cargo binding to IPO9 and cargo binding to importins generally (37).

The only study to broadly identify IPO9-specific cargo, conducted by Kimura et. al., used reconstituted nuclear import coupled to mass spectrometry and yielded 53 high-confidence and 254 medium-confidence cargo protein hits for IPO9 (14). While this approach works well for some of the better characterized importins, the overlap of identified cargo in this dataset with previously characterized IPO9 cargo was especially poor when compared to other importin•cargo interactions classified (Supplemental Figure 1A-D) and did not probe direct binding of the importin to cargo. This gap in knowledge limits our understanding of this essential biological process and thus, we set upon a holistic approach to identify IPO9 cargo and understand the key structural features that contribute to its selection for each of its cargo.

First, we robustly identify IPO9 cargo by reproducible enrichment with high-stringency cutoffs, recapitulating the majority of cargos identified in individual studies as well as 52 new cargos, and validate several of these with recombinant proteins. We then extend the use of hydroxy radical protein footprinting (HRPF) beyond purified recombinant factors to identify critical regions of IPO9 for cargo binding directly from the IP experiment, and discover that cytosolic IPO9 is predominantly occupied by cargo (38). From this, we leverage conservation, structure and phosphorylation data to predict critical regions for cargo binding and systematically perturb these elements to examine the consequences for the cohort of cellular cargo bound with quantitative proteomics. Through specific perturbations to the H7 loop, H8 loop and H18-19 loops of IPO9, we investigate how the unique non-HEAT repeat features of Importin 9 each contribute to binding specificity of cargo. The H8 and H18-19 loop specifically recruit cargo, while the H7, H8, and H18-19 loops promote binding of cognate cargo while preventing a class of secondary cargo that possess some affinity to IPO9 but do not bind to the WT importin. These experiments delineate the IPO9 features that govern which subsets of potential cargo predominate in a competitive paradigm for nuclear import. We find that RanGTP above cellular concentrations does not induce immediate competitive release of all cargo, rather, IPO9 cargos are variably susceptible to RanGTP binding, suggesting that additional accessory factors may be a general theme for nuclear cargo deposition. Overall, loop features explain a large fraction of IPO9 cargo binding and provide insight into the importin-specific features that mediate cognate binding to numerous diverse cargos via non-classical nuclear localization signals.

## Results

### Investigation into IPO9-bound proteins

To catalog a comprehensive list of cargo directly bound to Importin 9, we generated an inducible FLAG-tagged IPO9 cell line in a HEK293-FRT background (Figure 1A). In triplicate, we used mild digitonin lysis in the presence of WGA lectin to limit transport through nuclear pores (39) during lysis and fractionation of cells into cytoplasmic and nuclear samples. We then performed anti-FLAG immunoprecipitations for induced FLAG-IPO9 protein (Figure 1B). In the cytoplasmic IPO9 pulldown, we observe many proteins including dark bands on the lower end of the gel which likely correspond to the H2A-H2B histone dimer (Figure 1C), a canonical cargo (18, 20, 32, 36, 36). However, very limited bound cargos were detectable from the corresponding IPO9 immunoprecipitation from nuclear extracts, consistent with the idea that IPO9 is an importin, that is not involved in nuclear export (9). More importantly, the considerably smaller pool of IPO9 captured from the corresponding number of cells, is consistent with shorter nuclear than cytosolic residence time, and rapid cargo discharge upon nuclear entry resulting in limited steady-state cargo binding by the small fraction of IPO9 that is in the nucleus. Thus, we decided to perform mass spectrometry solely on the cytoplasmic fraction to determine protein cargos bound to IPO9.

**Figure 1.**
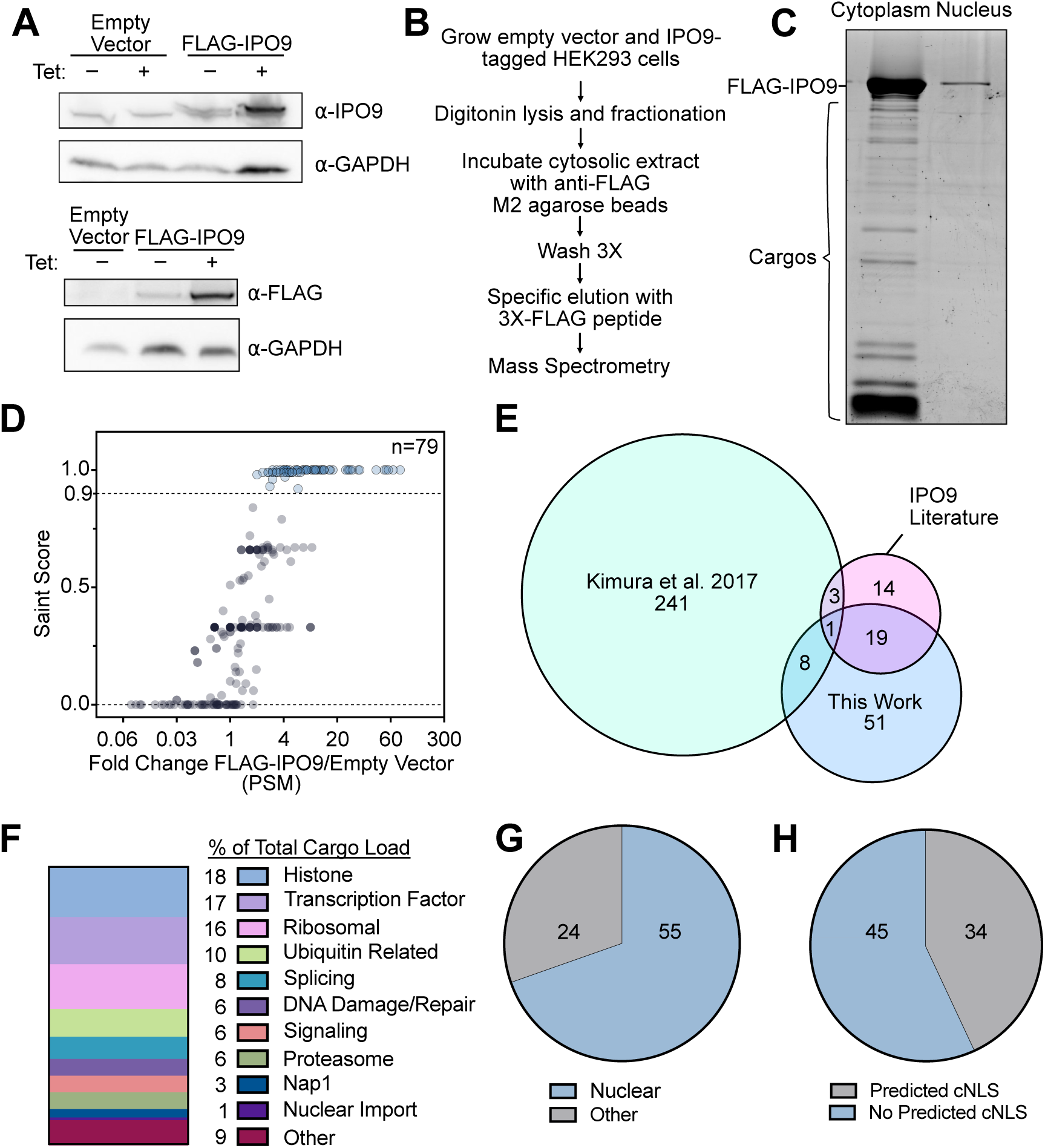
Identification of IPO9-bound proteome. **(A)** Western blotting confirms generation of FLAG-tagged IPO9 in HEK293 (T-Rex-293 FRT background with FLP-recombinase mediated site-specific integration). Expression with and without induction by tetracycline (Tet) shown, along with a blot for IPO9 for comparison to endogenous levels of protein; GAPDH blot serves as loading control. **(B)** Experimental workflow for pulldown of tagged importin 9 from HEK293 cell extracts. **(C)** SYPRO Ruby-stained 10% SDS-PAGE gel depicting 5% of the elution of cytoplasmic and nuclear FLAG-IPO9 immunoprecipitations for one replicate. **(D)** SAINT Express analysis of the mass spectrometry results of M2 pulldown from FLAG-IPO9 depicting fold-change of cargo protein versus control (empty vector cells) on the x-axis and SAINT score(40), used to calculate significance, on the y-axis. Seventy-nine cargo proteins detected with a SAINT score > 0.9. **(E)** Venn diagram showing the IPO9 cargo identified by individual cargo candidate-based studies. Previously investigated encompasses cargos identified via a variety of techniques *in vitro* and *in vivo* with IPO9 and/or its yeast homolog Kap114 (6, 9, 14, 18, 20–22, 31, 33, 74). **(F)** Total peptide counts for a given type of cargo relative as a fraction of total peptide counts of all cargos as an approximate measure of relative cargo bound. **(G)** Pie chart of Deeploc 2.1 proportion of cargo with nuclear localization annotation (45). **(H)** Pie chart depicting proportion of cargo predicted to have cNLS sequence by Deeploc 2.1.

With a stringent SAINT score (40) cutoff of 0.9, we identified 79 cargos as being significantly enriched in the FLAG-tagged IPO9 cell line relative to the control immunoprecipitations (IP) from parental HEK293-FRT cells that lack ectopically tagged IPO9 (empty vector control)(Figure 1D). There is excellent consistency between independent experimental replicate observations for each cargo and for experimental signal compared to background control (Supplemental File 1). We also identify many proteins unique to IPO9 versus control, some of which are not enriched enough for robust statistical determination, that may putatively be cargo of IPO9 (Supplemental File 1). We find 20 cargo proteins that have been previously identified by individual candidate-based studies and identify 51 novel cargos for IPO9 (Table 1). This overlap is better than measurements of other importins made by different groups (Supplemental Figure 1E) (14, 41, 42). Our overlap with individually studied cargos in the existing literature is much greater than Kimura dataset (14), despite having fewer protein hits (Figure 1E). Notably, we detect but do not find actin enriched over the empty vector IP, likely due to its abundant and sticky nature; it has been difficult to probe specific monomeric actin binding in pulldowns (35). We find SAINT >0.9 cargos are greatly enriched when compared to the whole cell mass spectrometry of HEK293 cells (Supplemental Table 1) (43), further supporting the specific nature of the IPO9 IP. Our method confirms direct binding by IPO9 to nine factors Kimura et al., noted to be imported into the nucleus in a reconstituted nuclear import system with IPO9 (histone variant H2A.Z, transcription factor NFYA, RNA methyltransferase NSUN5, splicing factors NUDT21/CPSF5, CPSF6, CPSF7, LUC7L, and RNA surveillance protein RBM26).

**Table 1.** Identification of IPO9 cargos. Cargo enriched over the control (untagged) sample SAINT score > 0.9. Cargos are listed in alphabetical order in two column sections depending on whether there is prior literature identifying a given protein as a cargo for IPO9. Type of evidence gathered for a given cargo are indicated below the table.

| Previously Identified |  | Novel Cargo |  |  |  |
| --- | --- | --- | --- | --- | --- |
| AKIRIN2 | PSMA4 | CALML5 | KLHDC2 | RIC8B | USP7 |
| *CPSF6* | PSMA7 | CCAR2 | KPNB1 | RPS5 | USP15 |
| *CPSF7* | PSMB1 | CCDC130 | MAGED2 | RPS10 | USP37 |
| *H2AFZ* | PSMB3 | CUL2 | MCM4 | RPS17 | YBX1 |
| **H2A** | PSMB4 | DDB1 | MCMBP | RPS18 | YWHAB |
| **H2B** | PSMB5 | FASN | MED25 | TCP1 | YWHAE |
| *LUC7L* | PSMD1 | FMC1 | MRPS7 | TEAD1 | YWHAG |
| **NAP1L1** | PSMD2 | GTF3C1 | OGT | TEAD4 | YWHAQ |
| **NAP1L4** | **RAN** | GTF3C3 | PCNT | TRIM28 | YWHAZ |
| *NFYA* | *RBM26* | H2AFX | PELO | TRIM35 | ZNF414 |
| *NSUN5* | *RPS7* | H2AFY | POGK | TRIP12 | ZNHIT6 |
| *NUDT21* | *RPS9* | HNRNPUL2 | PTOV1 | TUBA4A |  |
| PSMA1 | **TBP** | IRS4 | RANBP1 | UBC |  |
| PSMA2 | **TFIIB** |  |  |  |  |
| PSMA3 |  |  |  |  |  |

**Kap114 yeast homolog** *in vitro* direct binding *in vivo* indirect binding *in vivo* direct binding

We observe many major nuclear classes of proteins (Figure 1F) the proportions of which can be roughly ascertained by the abundance of a given class of proteins relative to total cargo protein peptide spectral matches (PSM). By this metric, the histone dimer represents about a fifth of total cargo bound to IPO9. Whereas novel transcription factor cargo TEAD1, TEAD4, GTF3C1, GTF3C3, in addition to TFIIB and TBP, collectively represent about 17% of the total bound protein to IPO9. We detect enriched AKIRIN2 and many core PSMA and PSMB proteasomal subunits (14). Signaling proteins from the 14-3-3 family (YWHAE, YWHAQ, YWHAB, YWHAG, YWHAZ) are also observed as IPO9 cargo. We note enrichment of the three main subunits of CFIm (44), an alternative splicing and polyadenylation complex, as Kimura did (NUDT21, CPSF6, CPSF7), indicating there may be co-import of the complex. NUDT21 binds to IPO9 directly with purified recombinant proteins (Supplemental Figure 1F), arguing that it mediates its own import and potentially tethers CPSF6 and CPSF7 to the importin. We also validate *in vitro* binding of ribosomal protein RPS18 as a new IPO9 cargo-it has been shown that RPS5 and RPS7 are bona fide IPO9 cargo, and our list generates many more ribosomal proteins that IPO9 binds and may import (Supplemental Figure 1F); ribosomal proteins are the second largest cargo category bound to IPO9.

Despite being immunoprecipitated from the cytoplasmic fraction, cargo candidates are largely predicted to be nuclear proteins by DeepLoc2.1 (Figure 1G) (45). DeepLoc2.1 (45) and NLSstradamus (46) also predict that the majority of cargo do not contain a cNLS (Figure 1H, Supplemental Figure 1G). We note that some cargo with annotated cNLSs do not necessarily use the cNLS for mediating import, as has been demonstrated for the H2A-H2B dimer (18). In exploring features of cargos for commonalities that may confer binding specificity, we see that many cargos are basic proteins (Supplemental Figure 1H-I). This is unsurprising for two reasons: many nuclear proteins that interact with acidic DNA and RNA biomolecules are basic, and IPO9 has an acidic, negatively charged inner surface that would accommodate binding and shielding of such charged cargos (33). Finally, we ran the 79 protein hits through peptide motif identification pipeline STREME (47) and found no significant enrichment for amino acid motifs, indicating that IPO9 cargos do not have a common identifiable linear nuclear localization signal (not shown). Since it is unclear what cargo-specific motifs may be dictating importin:cargo binding, we turned to IPO9 itself to investigate the different features on IPO9 that determine binding.

### Oxidative footprinting identifies key regions of IPO9 for cargo binding

To identify cargo binding regions of IPO9 in an impartial manner, we performed hydroxyl radical protein footprinting (HRPF) using Fenton chemistry on HEK293 FLAG-IPO9 pulldown samples in triplicate. While this approach has previously been used to query interaction surfaces *in vitro* with purified proteins (38, 48), this work is the first application of this benchtop HRPF strategy to complex immunoprecipitates (Figure 2A). Here, hydroxyl radicals are produced via Fe(II)-ethylenediaminetetraacetic acid (EDTA)-catalyzed Fenton chemistry on the lab bench without the need for specific instrumentation for hydroxy radical generation.

**Figure 2.**
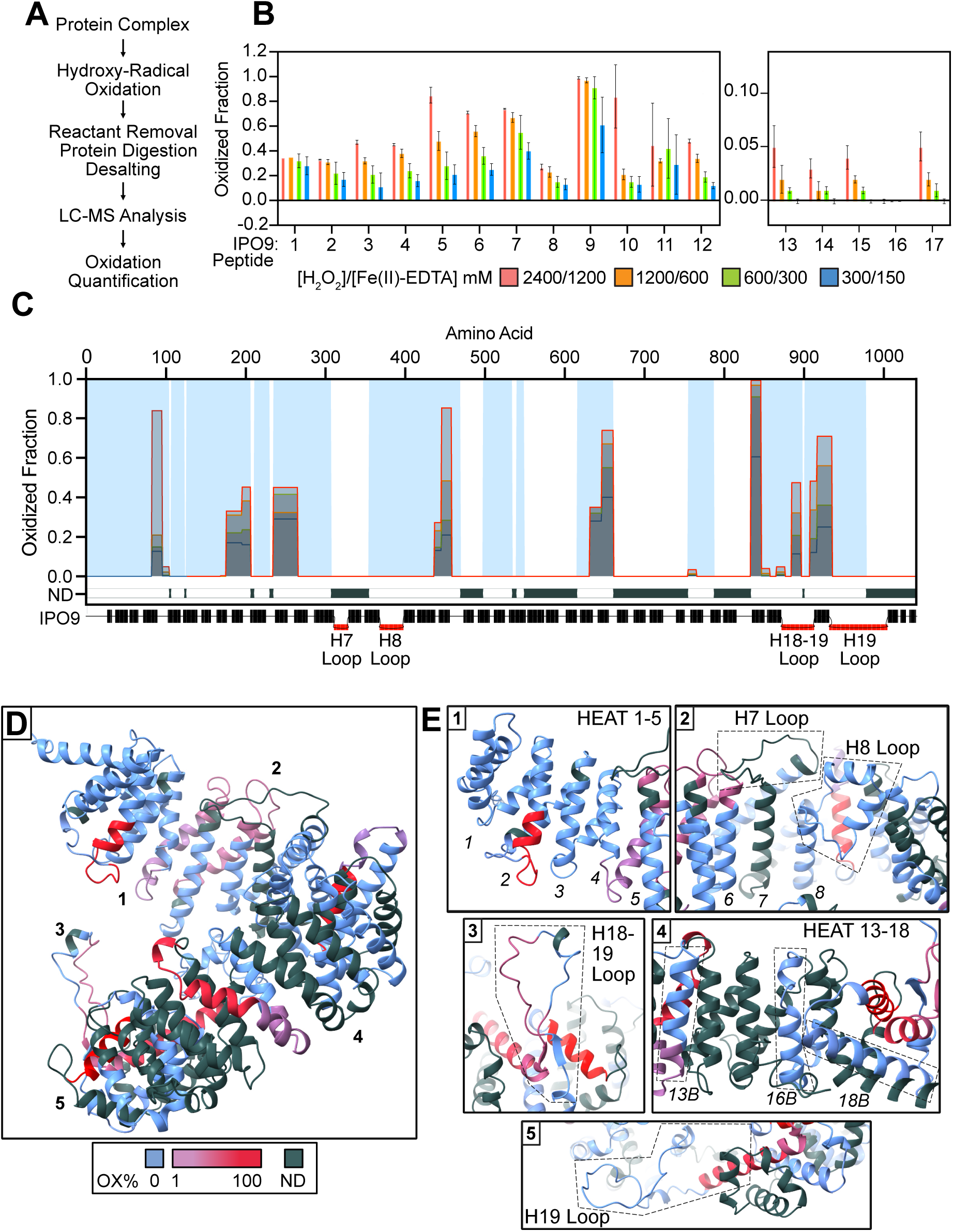
Oxidative Footprinting incriminates the internal surface and loops of IPO9 as cargo binding regions. (A) Workflow used for sample preparation in the oxidative footprinting (OX-FT) protocol. (B) Chart depicting peptides of IPO9 that were oxidized. Different colors correspond to concentrations of oxidant used, noted in the key below the chart. Peptide identities are noted in Supplemental File 2. (C) Chart of oxidation status (y-axis) mapped onto a linear depiction of the IPO9 peptide sequence (x-axis). Light blue shading corresponds to detected regions of IPO9, whereas ND corresponds to peptide regions of IPO9 that were not detected in the mass spectrometry. Grey areas correspond to oxidant concentration as noted in the chart in Figure 2B. (D) Predicted IPO9 structure is mapped with oxidation percentages onto the cartoon surface (75). Dark grey areas correspond to regions not detected via mass spectrometry, light blue equals detected but no oxidation, and the gradient from blue to red corresponds to increasing oxidation. (E) Zooming in on specific regions of the IPO9 structure depicted in (D).

In brief, FLAG-IPO9 pull down samples were treated with H_2_O_2_:Fe(II)-EDTA to induce oxidation on surface-accessible amino acid side chains on IPO9 (Figure 2A) and its bound cargo in a dose dependent manner. A clear dose-response curve is observed for a subset of peptides, indicating that these peptides are located on the surface of the protein (Figure 2B). Conversely, peptides where no oxidation is observed can be interpreted as either buried within the overall three-dimensional fold of the protein or bound to cargo. We do not normalize the percentage of oxidation according to the propensity of the respective amino acids to accept a hydroxy radical, therefore we do not interpret the magnitude of oxidation as the extent of surface accessibility, rather we use the presence of a dose dependent oxidation to infer that the amino acid is solvent accessible.

Between oxidized and unoxidized peptides, we obtain 65% sequence coverage of the IPO9 sequence (Figure 2C). The remaining 35% of the sequence is refractory to MS analysis after trypsin digest. Several IPO9 peptides exhibit clear dose response curves with small standard deviation across 3 biological replicates and 2 technical replicates (Figure 2B, Supplemental File 2). IPO9 Peptides 1, 10, and 11 lack a clear dose response, thus we are less confident in our assignment of these regions being on the surface. IPO9 peptides 13 to 17 show a relatively low magnitude of overall oxidation, nonetheless, we observe a dose dependent oxidation in this region suggesting they are solvent accessible.

When mapping the oxidation dose-response curves along the amino acid sequence of IPO9, we can observe specific regions that are oxidized readily (Figure 2C). On the contrary, there are many regions that are well detected but are not oxidized. The clear presence of both oxidized and unoxidized regions suggest the overall fold of the protein was not heavily perturbed before footprinting. It suggests that the IPO9, even when induced ectopically, is predominantly cargo-bound in the cytoplasm, and this apparent saturation survives the IP. Reinforcing this point, we obtain dose-response curves for some peptides of IPO9 cargos (Supplemental Figure 2A), but sequence coverage is not sufficient to make predictions about the location of binding. Interestingly, oxidation largely occurs throughout the HEAT repeats rather than the unstructured loops between them, supporting the hypothesis that the loops participate in cargo binding, as we detect the loops, but they are not oxidized. The exception is loop H7, as the tryptic peptide spanning this loop (amino acid 308-354) is not observable by mass spectrometry.

When the oxidation data is superimposed onto the apo-IPO9 structure (18), the inner surface is largely protected (HEAT repeats 1-4, 7, 8, 11, 13, and 16), whereas the lower and outer surfaces of IPO9 are oxidized and accessible, or not detected (Figure 2D-E). Surprisingly, we find that the tertiary disordered loops (H8, H18-19, H19), despite being flexible and solvent-accessible in the apo structure, are either not oxidized at all or less than half of them carrying an oxidation modification, indicating they may be participating in cargo binding. Unfortunately, we do not have MS sequence coverage for the H7 loop (H7) along with Heat Repeats 10, 12, 15, 17, and 20 and are unable to comment upon their surface accessibility.

In an attempt to generate an apo-IPO9 control, we stringently purified recombinant IPO9 and performed the HRPF protocol. Surprisingly, we observe a similar oxidation profile to the mammalian extract IPO9, seemingly due to widespread copurification of basic bacterial ribosomal and nucleic acid binding proteins (Supplemental Figure 2B-C). These data hint that IPO9 prefers to be in a cargo-bound state, which remarkably survive stringent purification protocols.

### Conservation analysis and PTMs implicate disordered loops as regions of interest

Despite gross structural similarity, importins are thought to have distinct niches of cargo from one another (9, 14, 42). Several features vary among importins, such as the pitch, radius, number of HEAT repeats and inter-HEAT repeat loop regions–stretches of amino acids between heat repeats that vary in length and may be structured or unstructured (Supplemental Figure 3A-B). Lack of evolutionary conservation within the karyopherin family defines unique features amongst importins that may provide specificity for respective cargo niches. Many importins have a disordered inter-HEAT stretch around residues 300-400 that is used for Ran binding (49–53), though the size varies greatly (Figure 3A, Supplemental Figure 3A) (54). IPO7, IPO8, and IPO11 also have large C-terminal loops, though their exact size and position vary relative to IPO9’s H18-19 and H19 loops. Conversely, regions that are highly conserved among a given importin’s orthologs point to specific features that may be critical for proper protein function. Conservation analysis of IPO9 homologs through CONSURF, mapped onto the sequence of IPO9, shows that the inner surface of IPO9 is largely conserved, as are portions of the H7, H8 and H18-19 loops (Figure 3B, Supplemental Figure 3C) (55–60). Another indicator that loops play key discriminating roles in cargo affinity was the presence of post-translational modifications (PTMs). We found phosphorylation sites at serine and tyrosine residues on all the loops of IPO9 (Figure 3C-D). Strikingly, the S322 residue on the H7 loop is found to be phosphorylated in over 78% of PSMs and the S947 residue on the H19 loop is phosphorylated on about 60% of PSMs. Functionally, this pervasive phosphorylation would make already acidic loops even more charged and could serve to modulate bound-cargo distributions.

**Figure 3.**
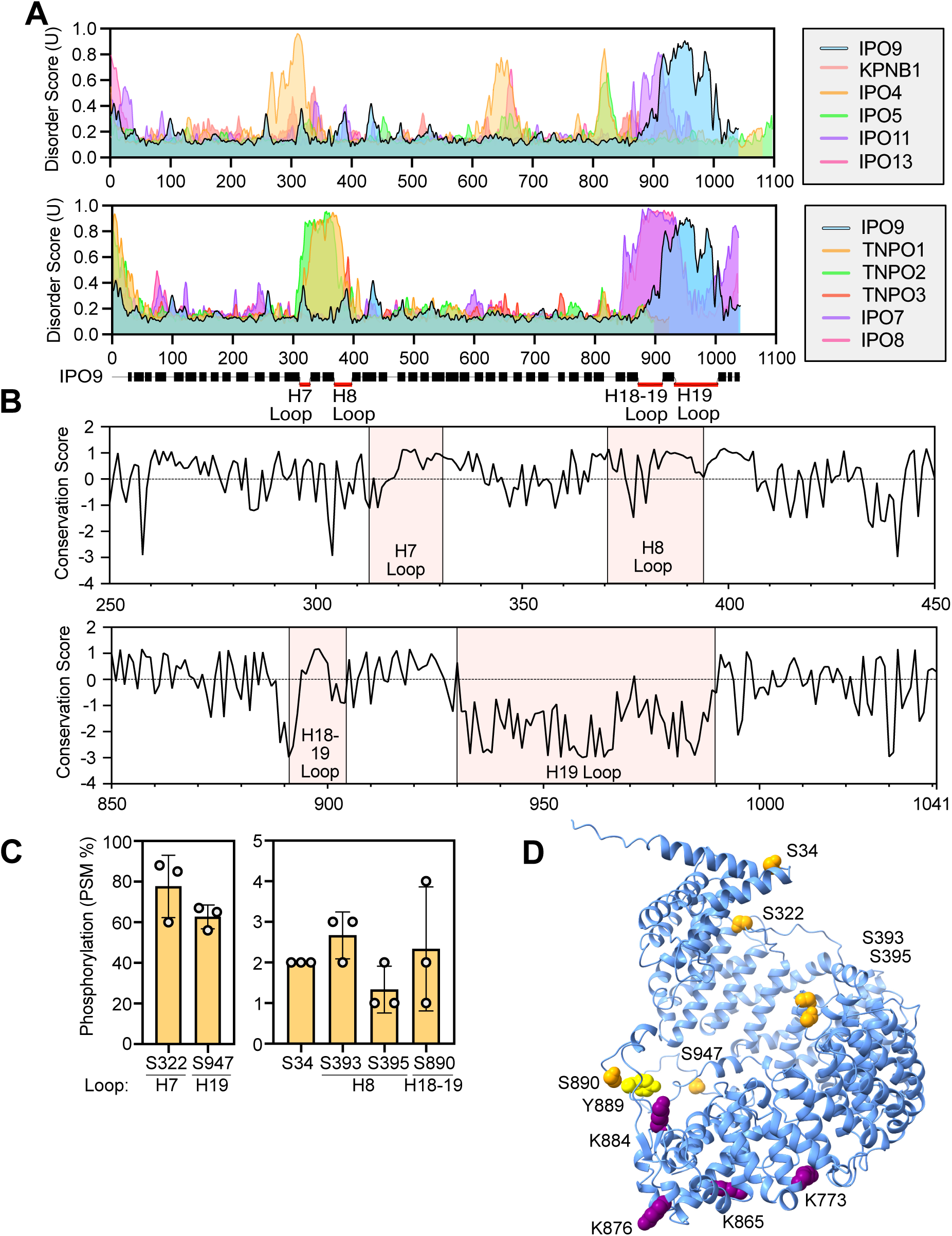
Conservation and PTM analysis implicate the loops of IPO9 as unique features that differentiate IPO9 from other importins. (A) Alignment of disorder plots of the importins superimposed on top of one another. Y-axis scale is from .15-1.0 disorder to look at solely the more disordered sections of the proteins. Coloring of each protein noted on the right (76). (B) Conservation of regions of IPO9 protein with loops present. Average conservation equals 0 (dotted line). The parts of the H7, H8 and H18-19 loops are highly conserved, while the H19 loop is not. Computed via CONSURF-DB (77) (C) Graph of phosphorylation-site analysis of IPO9 pulldown data. (D) Post-translational modifications that have been found by us or other groups (n>1) are imposed onto the structure of IPO9 (61).

### Comprehensive dissection of the inter-HEAT loop features of IPO9

Considering this evidence pointing towards the loops of IPO9 as modifiable, cargo-bound regions, we decided to deeply probe contributions of these loops of IPO9 to cargo binding. We generated FLP-FRT HEK293 lines of these mutants and blotted to confirm commensurate protein expression and apparent stability (Supplemental Figure 4A-B) in a background containing two endogenous WT IPO9 alleles. This ectopic expression format, rather than a knock-in that alters the endogenous allele, has multiple benefits: 1.) reduced risk of toxicity, given IPO9 is an essential protein; 2.) ease of line construction; and 3.) ensuring equivalent expression level across the cohort of ectopic protein through site-specific integration and selection (absent differential protein stability). However, despite several attempts, we were unable to generate H19 loop deletion lines with detectable protein expression; perhaps this mutation renders the protein unstable or cells that express the mutant die due to a dominant negative effect of the mutant. Given the low sequence conservation and negative charge over two D/E regions of the loop, it was unclear what exact regions to target with point mutations in this region. In the next sections, we assess consequences for cargo binding of H18-19 point mutations and deletions of the H7 and H8 loops through IP-MS of the IPO9 mutants.

### Exploring the cargo binding contributions of the H18-19 loop through point mutations

Although the H18-19 loop has been shown to be critical for H2A-H2B histone dimer binding in vitro (18), whether it is similarly consequential for other cargo binding remains uncertain. To test the generality of this recognition mechanism, we created a charge swap mutation, R898E, of the arginine that forms a salt bridge with the histone dimer acidic patch in the crystal structure (PDB:6N1Z) (Figure 4A)(18). Consistent with prior *in vitro* findings, SDS-PAGE of the IP elution indicates that histone dimer binding is absent (Figure 4B). Remarkably, through IP-MS, we find the histone dimer proteins H2B and H2A (including variants) are the only proteins that are significantly depleted in the mutant as compared to the WT IPO9 IP, consistent with this being an energetically essential component of recognition solely for histone cargos (Figure 4C). We define cargos that are present in the WT IPO9 IP as cognate cargos– many cognate cargos are present in R898E mutant, and a subset are specifically enriched (SAINT > 0.9) in the mutant (Figure 4D). Specifically, we note increases in many ribosomal proteins, splicing factors and transcription factors (NFYA, YBX1, YBX3) by PSM. We also observe a cohort of “secondary cargo”, which we define as cargo proteins not detected in the WT IP but observed in the mutant. Among these secondary cargos we see more ribosomal proteins, translation factors, and other nuclear proteins.

**Figure 4.**
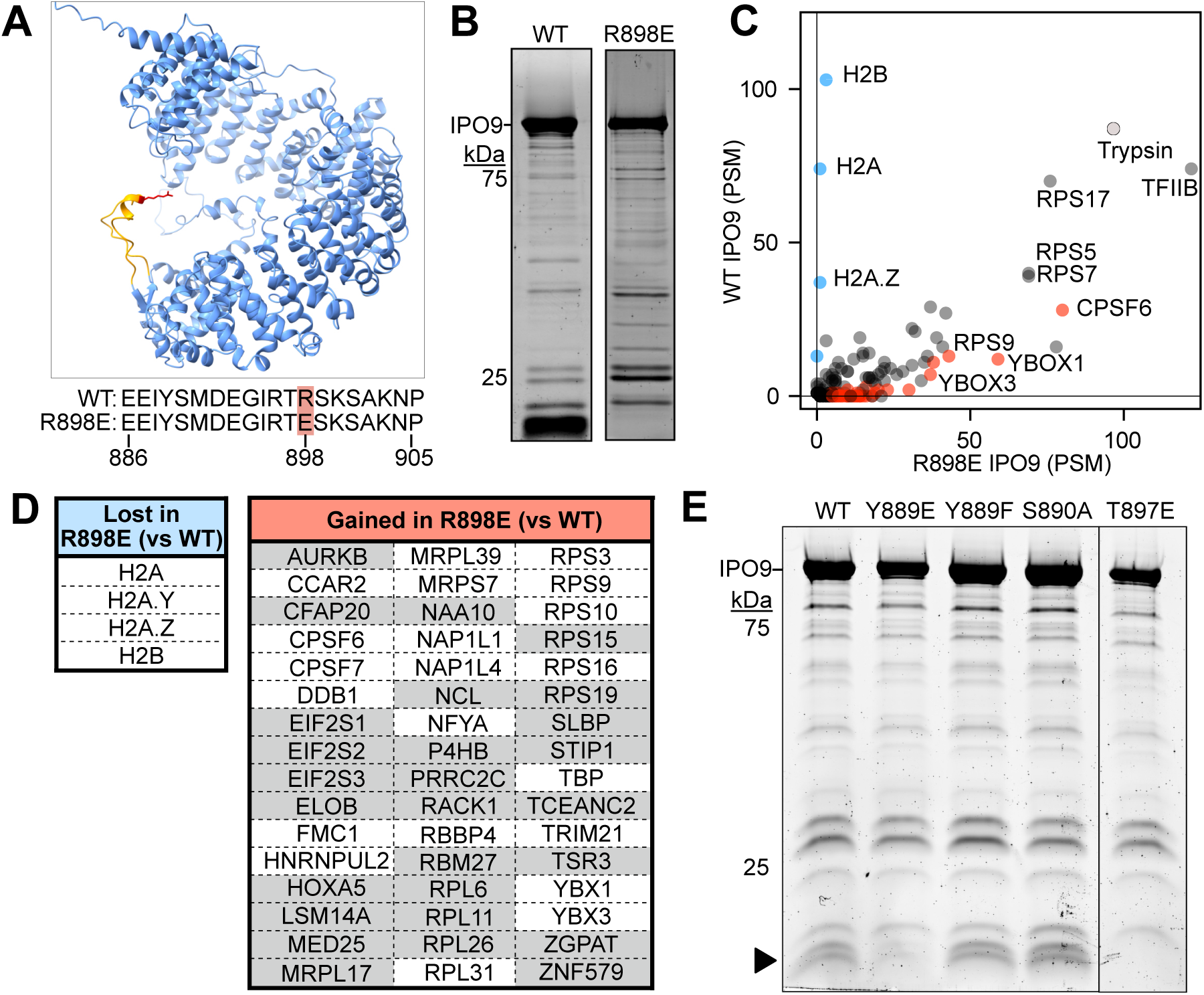
The H18-19 loop is critical for histone dimer binding and histones competition shapes the cargo binding profile of IPO9. (A) Structural representation showing H18-19 loop on IPO9 Alphafold structure. (B) Sypro Ruby-stained 10% SDS-PAGE with 5% of WT and R898E mutant elution loaded. (C) Graph depicting protein spectral matches (PSM) of WT versus R898E mutant. Dashed line depicts theoretical 1:1 binding. Exogenous bovine trypsin (light grey dot) serves as an internal control for equivalent mass spectrometry sample preparation and loading. SAINT > 0.9 samples enriched for WT (blue) and 898E (red) highlighted on graph. (D) Table showing the SAINT > 0.9 enrich cargos for WT versus the R898E mutant and *vice versa*. Cargos shaded in grey are new, secondary cargos that are not found in WT IPO9 IP samples. (E) Sypro-Ruby stained 10% SDS-PAGE gel of small-scale IPO9 mutant immunoprecipitations.

To further interrogate the function of the H18-19 loop, we turned to other features present on the loop. Phosphorylations within this loop occur at residues T897, S890 and Y889 (61). To test whether the charge effect of H18-19 loop phosphorylation mirrors the effect of the R898E mutation, we made phospho-mimetic and phospho-null point mutations on the loop in other locations where phosphorylation events take place and did a smaller scale IPO9 IP (Figure 4E). Indeed, by SDS-PAGE we see loss of H2A-H2B dimer binding in the phospho-mimetic mutants (Y889E, T897E) but not in the phospho-null mutants (Y889F, S890A), indicating that these loop phosphorylations could regulate H2A-H2B binding and import.

Overall, we demonstrate through multiple point mutations that the H18-19 loop is essential for *in cellulo* histone dimer binding, in alignment with previous structural studies (18, 36). The specific response, where we only lose binding of the H2A-H2B dimer, gives us confidence in the robustness and reproducibility of the technique in that the changes we see in IP-MS signal are reflective of both altered biochemical affinity that the mutant importin has for a given cargo. Next, we applied this approach to investigate the cargo binding contributions of the H8 loop of IPO9.

### Changes in cargo binding upon H8 loop deletion

The H8 loop sits on the upper-middle inner surface the IPO9 solenoid (Figure 5A) and has two conserved regions that are negatively charged (Supplemental Figure 5A). A cryo-EM structure and hydrogen-deuterium exchange coupled to mass spectrometry indicated that, in addition to the N-terminal HEAT repeats of IPO9, H8 loop residues 373-385 are bound (36, 62). However, it is unclear what contributions that this loop makes to cargo binding, nor precisely the extent to which RanGTP binding may antagonize cargo engagement. We chose to perturb this loop through a deletion modeled after the yeast Kap114 RanGTP binding mutants (63) (Supplemental Figure 5B)– amino acids flanking this deletion are polar and should afford sufficient linker length and flexibility to span the intra-HEAT gap.

**Figure 5.**
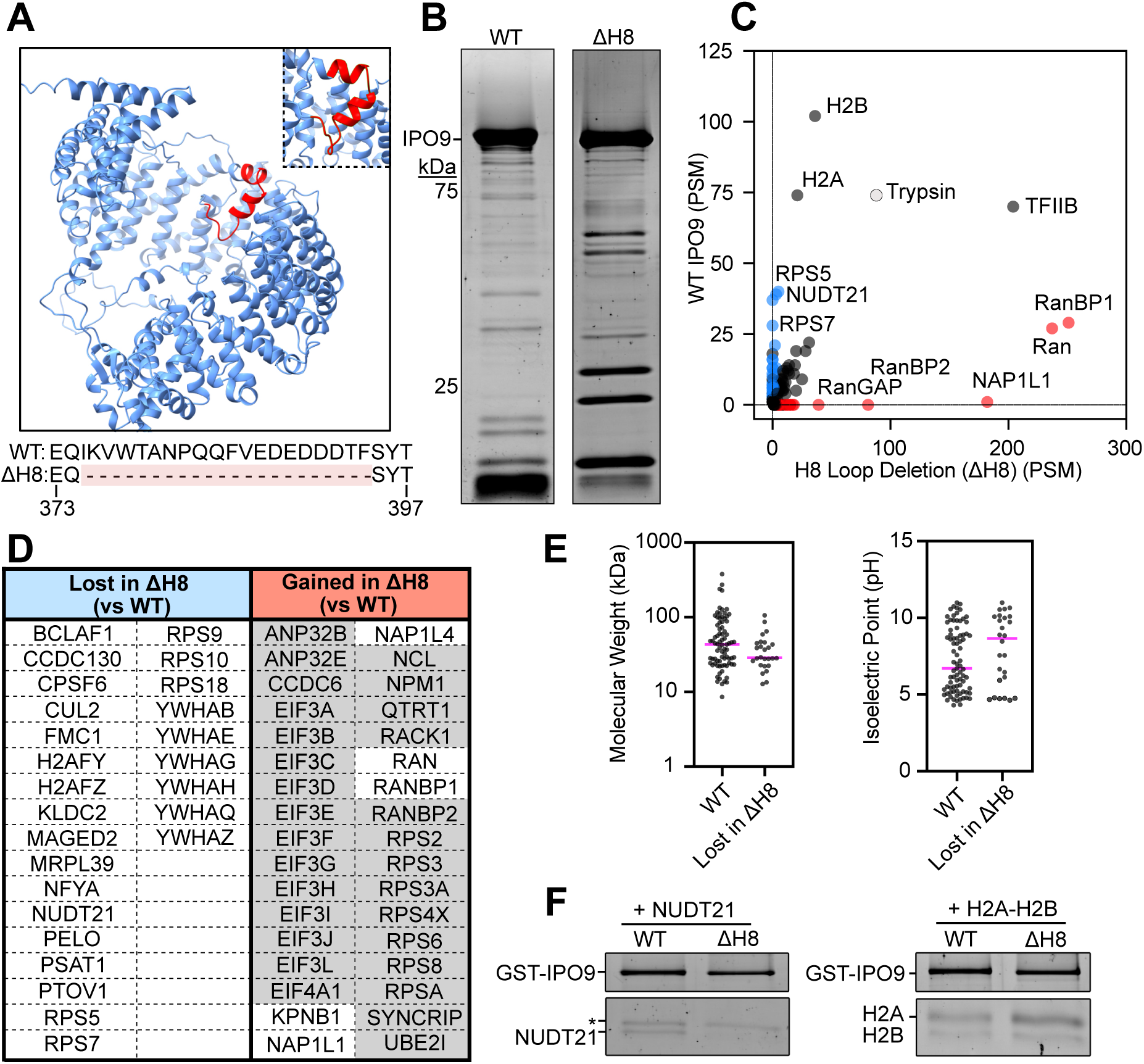
The H8 loop serves to both recruit primary cargo and dispel binding of secondary cargo. (A) Structural representation of H8 loop deletion on IPO9 Alphafold structure with peptide sequence and deletion made depicted below. (B) Sypro Ruby-stained 10% SDS-PAGE with 5% of WT and H8 deletion mutant elution loaded. (C) PSM comparison of wild type (WT) versus H8 loop deletion (ΔH8). Line depicts theoretical equivalent binding between WT and mutant defined by IPO9 capture. Trypsin is shown to show internal control for mass spectrometry sample preparation. (D) Table showing (left) cargos enriched in the WT sample over the ΔH8 samples and (right) cargos enriched in the ΔH8 IPO9 sample over wild type sample with SAINT score >0.9. Cargo with grey background represent secondary cargo that are not present in the wild type IPO9 sample. (E) Comparisons of molecular weight and isoelectric point of cargos lost in the H8 loop deletion to WT cargos. Computed by EXPASY calculator. (F) GST-IPO9 pulldowns (WT and ΔH8 Loop mutant) with histone dimer cargo and NUDT21.

The pulldown gel shows both specific enrichment and specific loss of various cargos (Figure 5B), indicating that the loop is critical for defining the cohort of IPO9 cargos. Through mass spectrometry, we observe complete loss of proteins classes that likely directly bind the H8 loop (Figure 5C-D), including some ribosomal proteins (RPS5, RPS7, RPS9, RPS10, and RPS18), 14-3-3 proteins (YWHAE, YWHAQ, YWHAB, YWHAH, YWHAG, and CUL2) and histone variants H2A.Y and H2A.Z. Consistent with H8 loop charge complementarity within the IPO9 central cavity playing a key role in binding their energetics, cargos lost are smaller than average cargo bound in WT case and tend to be more positively charged (Figure 5E). We note loss of transcription factor NFYA and splicing factors CCDC130 and the CFIm complex (NUDT21 and CPSF6; CPSF7 is also lost but not detected at high enough levels for statistical enrichment analysis). In an IP with recombinant purified factors, we find that NUDT21 requires the H8 loop to bind to IPO9, while canonical H2A-H2B dimer binding is unaffected (Figure 5F, Supplemental Figure 5C). In the absence of the H8 loop, we find recruitment of a new secondary set of proteins – cargo that have some affinity for IPO9 but are either outcompeted by cognate cargo or normally excluded by the intact H8 loop. We see gain of many more ribosomal proteins of the small subunit. The gain of some ribosomal binding proteins with concomitant loss of others indicates ribosomal protein association is not merely a charge-based interaction, but that sequence or structural components of specificity are at play regulating binding affinity of basic proteins.

Closer examination of the cognate cargos that increase in peptide counts upon H8 loop deletion, reveals large increases in Ran, RanGAP, RanBP1, RanBP2, and Nap1. The large increase in Ran engagement is surprising as this loop deletion was initially designed to be a Ran-binding deficient mutant based on yeast Ran binding mutant data and the seeming importance of this interface deduced from the RanGTP-IPO9 structure (Supplemental Figure 5B)(62, 64). Our mutant does not delete residues 393-395 which are absent in the Kap114 mutant and perhaps this is what induces the dominant Ran-binding phenotype, or the RanGTP contact surface outside of this loop is energetically more important. RanGAP and RanBP1 associate with the cytoplasmic fibrils of the nuclear pore, suggesting that the mutated H8 loop may increase IPO9-pore binding. The association of histone chaperone Nap1 with IPO9 without corresponding increase of the histone dimer is somewhat surprising given previous literature exploring their binding in tandem (18, 20, 32, 36, 62).

### The H7 Loop discriminates against secondary cargo binding

For our final IPO9 perturbation, we turned to the H7 loop, which has not been previously probed or implicated in cargo or RanGTP binding in any manner to our knowledge. When we delete the conserved portion of the H7 loop with a two amino acid linker (Figure 6A), gel analysis of the IP shows no obvious loss of cargo but gain of bands (Figure 6B). In the IP-MS, we find no specific loss of protein cargo, indicating that the H7 loop does not impart much favorable binding enthalpy to IPO9 cargo. Rather, we observe that deletion of the loop results in the significant binding of many new cargos (Figure 6C, 6D), consistent with it restricting potential cargo binding by repulsion. Intriguingly, the increase in binding is almost entirely due to addition of secondary cargo, mainly small and large ribosomal subunit proteins and translation factors. It is possible that the loss of the H7 loop causes IPO9 to lose some ability to discriminate amongst these highly basic charged proteins, potentially leading to their inappropriate nuclear import. Absence of this gatekeeping function of the loop could increase association of IPO9 with mature ribosome and translation factor complexes (eIF cohort of proteins). This result for the H7 loop reinforces that structural features of IPO9 gatekeep cargo binding to prevent spurious import.

**Figure 6.**
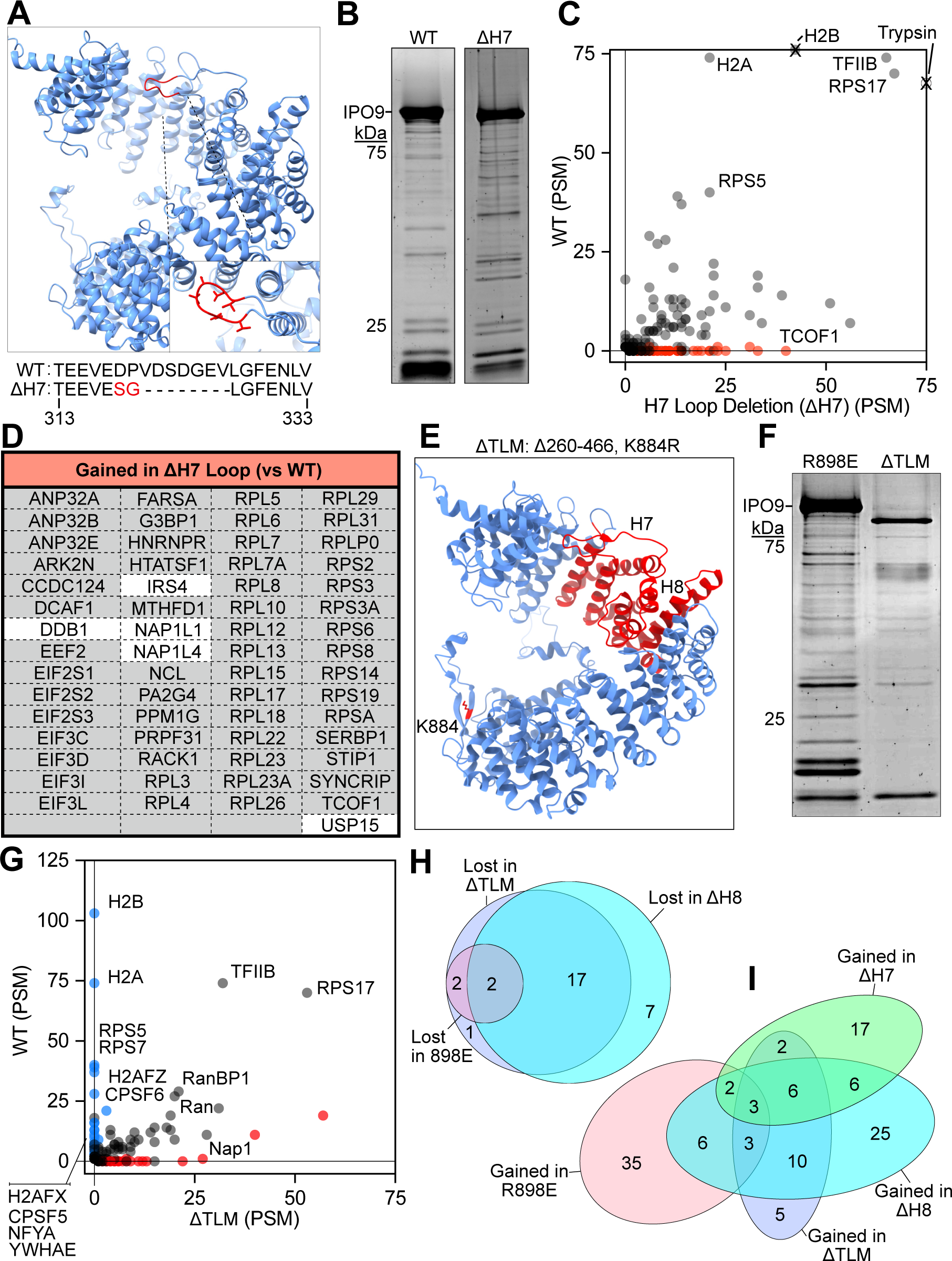
The H7 loop of IPO9 is primarily guards against secondary cargo binding and the ΔTLM mutant recapitulates individual loop perturbations. (A) Structural representation of H7 loop deletion on IPO9 Alphafold structure with peptide sequence and deletion made depicted below. (B) Sypro Ruby-stained 10% SDS-PAGE with 5% of WT and H8 deletion mutant elution loaded. (C) Graph with wild type (WT) versus H7 loop deletion (ΔH8). Line depicts theoretical equivalent binding between WT and mutant. Trypsin shown to show internal control for mass spectrometry sample preparation. (D) Table showing cargos enriched in the ΔH7 IPO9 sample over wild type sample with SAINT score >0.9. Cargo with grey background represent secondary cargo that are not present in the wild type IPO9 sample. (F) Structural representation showing the ΔTLM mutant. (G) Sypro-stained 10% SDS-PAGE gel showing FLAG immunoprecipitations of the R898E and ΔTLM IPO9 mutants. (H) Graph of the WT versus ΔTLM immunoprecipitations. (I) Venn Diagram of overlap of cognate cargos lost in ΔTLM vs the R898E and ΔH8 mutants. (J) Venn Diagram of secondary cargos gained in ΔTLM vs R898E, ΔH7 and ΔH8 mutants.

### Large structural perturbation validates critical loop regions for cargo binding while retaining binding to a subset of cognate cargos

To validate contributions of the loops to IPO9 cargo binding and test the limits of a retained cognate cargo recognition, we generated a mutant that deletes a central portion of IPO9, including the H7 and H8 loops, and has the H18-19 loop perturbed (Δ260-466, K884R, Triple Loop and Middle mutant (ΔTLM)) (Figure 6E, Supplemental Figure 6A). Despite lower apparent expression, potentially due to stability defects, we detect some full-length IPO9ΔTLM and verify that many cargos are still bound to this severe mutant via SDS-PAGE (Figure 6F). Through mass spectrometry we find that this mutant is still able to bind many cognate cargos, including Ran, RPS17, the proteasome, and Nap1 (Figure 6G, Supplemental Figure 6B-C). IPO9ΔTLM fails to bind cognate cargos (the H2A-H2B histone dimer, CPSF5 and CPSF6, 14-3-3 proteins, etc.) that are also lost in the R898E and ΔH8 mutants (Figure 6H, Supplemental Figure 6C). This mutant also significantly enriches many secondary cargos that are found in the ΔH8, ΔH7 and R898E mutants (Figure 6I). IPO9ΔTLM confirms that the H8 loop and surrounding residues are not necessary for Ran binding *in cellulo.* However, we note that the ΔH8 mutation alone markedly increases Ran binding in conjunction with RanBP1 so the H8 loop may play a role in modulation of Ran release.

### Most IPO9 cargos are remarkably insensitive to RanGTP release

We were intrigued that the large increase in Ran binding in the H8 loop deletion did not prevent IPO9 from binding to many cognate and secondary protein cargos. We also wanted to determine whether the loss of cargo resulting from the H8 loop deletion was a function of the large increase in Ran and RanBP1 binding as binding of the Ran release factor may limit the binding of Ran-sensitive cargo to IPO9. To test the Ran-sensitivity of IPO9 cargos, we performed a pulldown from HEK293 cells on FLAG-tagged WT IPO9 with attendant washes but prior to 3X-FLAG peptide elution we incubated the resin-bound IPO9 complexes with a 10µM recombinant RanGTP (Figure 7A), roughly double the physiological concentration of RanGTP in the nucleus (4.3µM) (65). We observe widely ranging RanGTP sensitivity of IPO9 cargo (Figure 7B). In good accordance with previous literature (18), we find a cargo that has been shown to be very insensitive to solely RanGTP-mediated release, the H2A-H2B dimer, falls at around 0.2-fold enrichment in the RanGTP versus FLAG elution samples. The proteasome, but not its IPO9 adaptor AKIRIN2, is sensitive to RanGTP binding, in concurrence with recent findings (17). NFYA, RPS17, RPS9, TFIIB and TBP are similarly insensitive to RanGTP binding as the histone dimer which indicates they may require additional factors for release once inside the nucleus (Supplemental Table 4). Conversely, CPSF6, NUDT21, YBOX1 and YBOX3 are sensitive to RanGTP binding. In comparison to proteins lost in the H8 loop deletion, we find decreased overall sensitivity to RanGTP-wash than the total population, indicating that RanGTP binding is likely not responsible for loss of most of these proteins in the H8 loop deletion, rather the H8 loop itself is being used to specifically recruit these proteins (p=.0025, two-sided t-test) (Figure 7B). We also observe some secondary cargos in this round of mass spectrometry--there is greater depth due fractionation over two runs with weaker binding cargo in the wash. We detect about half of all secondary cargos for each mutant--detected cargo as a whole are more Ran-sensitive than cognate IPO9 cargos (p=.0002, two-sided t-test). The ΔTLM secondary cargos are less RanGTP sensitive than entire group of secondary cargos (p=.0174, two-sided t-test), perhaps indicating increased affinity for IPO9 and less effective competition by RanGTP.

**Figure 7.**
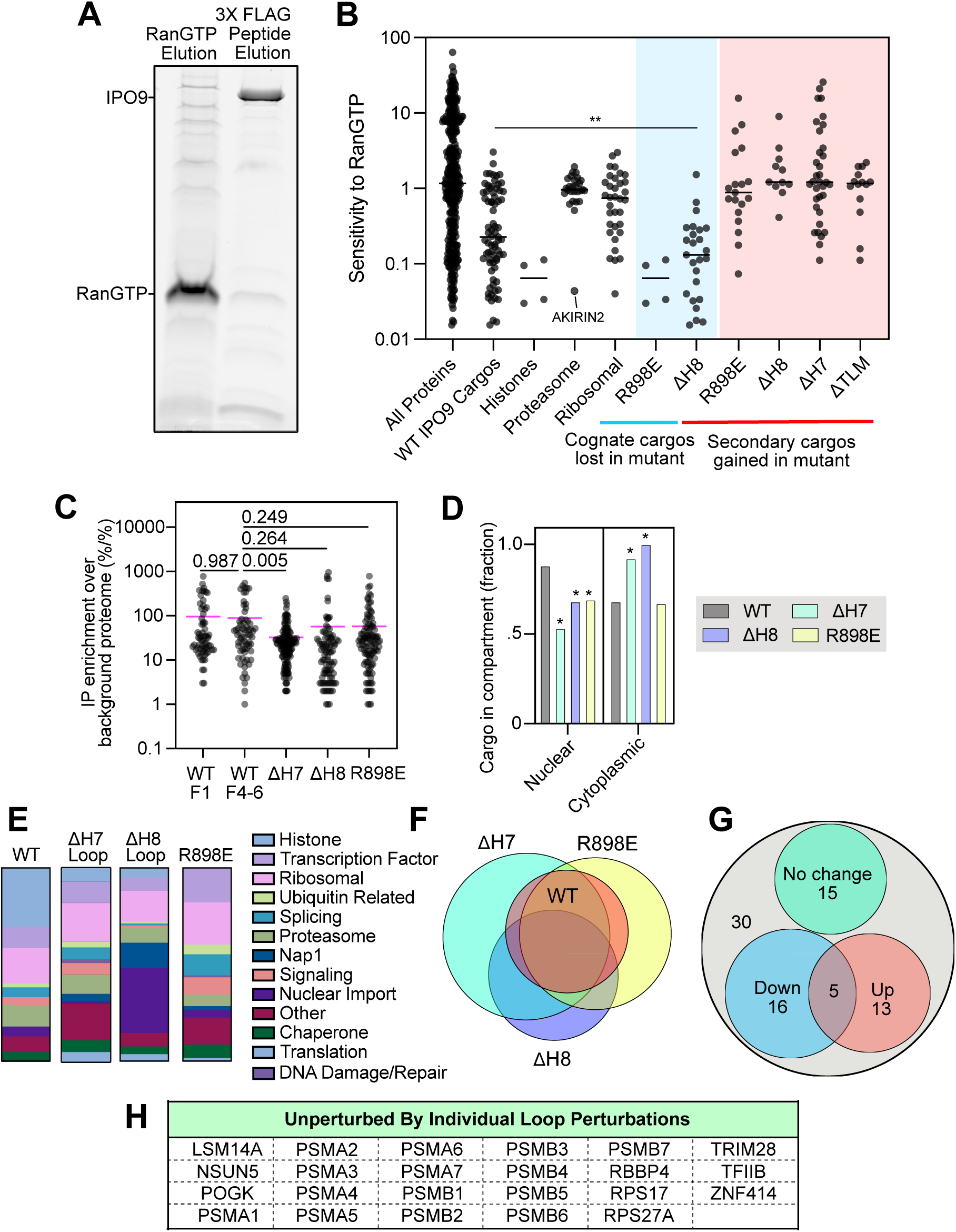
IPO9 cargos are variably sensitive to RanGTP; Loop perturbations modulate cargo binding to IPO9. (A) Sypro-stained 10% SDS-PAGE of 5% elution for a FLAG-IPO9 pulldown followed by a sequential RanGTP elution then 3X-FLAG peptide elution. (B) Graph showing the variable sensitivity of IPO9 cargo to RanGTP, with the y-axis corresponding to increasing sensitivity to RanGTP. Two-sided student’s t-test used to determine *p* values of a comparison of RanGTP sensitivity of WT cargos to those lost in the H8 loop deletion (SAINT>0.9). (C) Relative enrichment of cargos bound to ectopic constructs vs cargo abundance in HEK293 cells as reported by (43). F1 refers to Figure 1 IPO9 IP. F4-6 refers to WT IPO9 cargo IP used in comparison to IPO9 mutants. (D) Nuclear and cytoplasmic percentages of cargo for WT and mutants. (E) Cargo load categories of WT IPO9 versus mutants. (F) Venn diagram depicting quantitative overlap of all cargo detected in WT samples versus each mutant sample. (G) List of cargo that were unchanged through IPO9 loop perturbations. (H) Venn diagram showing cognate cargo that were measurable unchanged (green), increased (red) or decreased (blue) by the IPO9 loop perturbations. Cargos that were not abundant enough in mutants to determine if significant changes occurred in (grey).

Given that the histone dimer is released from IPO9 in concert with nucleosome-bound DNA (18), we were curious whether IPO9 itself has affinity for nucleic acids and if this could be an additional factor aiding in cargo release. Many of IPO9 cargo are nucleic acid binders (Supplemental Table 1) and IPO9 has been posited to bind the 5’ UTR of interferon epsilon mRNA, however this evidence was indirect (25). In a fluorescence polarization direct RNA binding assay using recombinant IPO9, we find no detectable affinity between IPO9 and WT or hairpin-loop mutated interferon epsilon RNAs (Supplemental Figure 6D). This does not rule out that nucleic acid could bind a cargo itself and effectively chaperone a surface of the cargo to promote release, as has been found for H2A-H2B (18).

The lower-than-expected sensitivity of cargo to prolonged incubation with high concentrations of RanGTP indicates secondary nuclear release factors may be a general requirement for many IPO9 cargos. We find that the H8 loop of IPO9 is a critical determinant to recruit and repel cargos while also facilitating RanGTP release factor binding.

## Discussion

Hundreds of proteins must be specifically transported from the cytoplasm, where proteins are synthesized, into the nucleus, where they govern all aspects of nuclear structure and function. Between the H2A-H2B dimer, TFIIB, TBP, the proteasome, and actin, IPO9 has demonstrated an ability to bind and transport a diverse array of protein cargos into the nucleus (18, 21, 22, 31, 35). Here, we carry out a large-scale mass spectrometry-based approach to catalog cargos directly bound to IPO9, recapitulating binding of cargos identified in foundational yeast literature in a mammalian context (20, 21, 31, 66) and finding many novel cargos. These cargos largely do not contain classical nuclear localization signals, nor any identifiable linear signal peptide sequence. Our HRPF data indicates that the inner core of IPO9 is largely occupied by cargo and that the loops between HEAT repeats are also occupied by cargo. This lack of a substantial population of unbound IPO9, even when it is slightly overexpressed, argues that all cargo binding is a competitive process, determined by the composite of each cargo’s affinity to the importin and its cytosolic concentration. Evolutionary analyses and post-translational modification data further suggest that these inter-HEAT repeat loops of IPO9 are conserved and subject to a number of post-translational modifications We use an *in cellulo* biochemical structure-function approach to define the loop contributions to cargo selection This approach and accompanying experiments, permits a view into the complex competitive binding equilibrium of cargo and enables an understanding of the redistribution of bound cargos and concomitant exclusion of inappropriate cargo.

Through targeted loop perturbations, we show disrupting the H18-19 loop via charge switch of the R898 residue markedly disrupts solely histone cargo binding. We dissect the role of the H8 loop and find that it specifically promotes the binding of a class of IPO9 cargo and decreases binding of secondary cargo, some of which may not be appropriate for nuclear import. We find that the H7 loop, while not recruiting specific cargo, excludes a large class of secondary cargos. We show that the sensitivity of IPO9 cargos to prolonged incubations in high concentrations of the RanGTP release factor is highly variable, and argue that additional release factors for IPO9 cargo in the nucleus may go beyond previously identified cargos H2A-H2B, AKIRIN2 and TFIIB (17, 18, 31). Cargos sensitive to loop perturbation provide a framework for future researchers to understand how IPO9 is interacting with its cargo and a means to disrupt these interactions. For example, our data argue that H2A/-2B import can be selectively disrupted by R898E mutations without impact to other cargo import by IPO9. This raises the prospect that fine-grained mutations to the identified loops could selectively impair import of other cargos. Detailed studies of IPO9 reveal broader principles of importin/exportin-cargo selection and prompts similar studies investigating the mechanisms governing how any nuclear protein is transported.

### Oxidative Footprinting of native IPO9 complexes

HRPF supports a model in which IPO9 is largely cargo bound, as oxidation is primarily observed within HEAT repeats rather than the unstructured loop regions. The clear separation of oxidized and protected regions suggests that the overall fold of the protein and cargo remain stable and not perturbed during immunoprecipitation and subsequent footprinting. Notably, regions detected by mass spectrometry yet remain unoxidized, specifically the loop regions, including H8, H18-19, and H19, provide strong evidence for involvement in cargo binding, as their protection cannot be attributed to lack of detection or propensity for the amino acids to accept a hydroxy radical. This is further reinforced by the preferential oxidation of residues within HEAT repeats, indicating these regions are largely solvent-exposed and unlikely to be a major contributor to cargo binding. Importantly, the application of benchtop HRPF to affinity purified (AP) complexes represents an additional dimension of information obtained from AP-MS experiments. This strategy enables simultaneous identification of bound cargo and insight into the binding interface on the bait protein and possible also on the interacting partners. However, incomplete sequence coverage, especially for the baits limit our conclusions. For other complexes that are less variable in their composition, this HPRF approach with AP-MS material holds even greater promise. Beyond limited bait coverage, IPO9 regions with sparse MS evidence, such as the H7 loop, peptides 1, 10, and 11 and several HEAT repeats, also remain ambiguous. Despite these limitations, the footprinting data supports a model in which IPO9 engages cargo through a combination of inner surface contacts and flexible solvent-exposed loops, consistent with multiple different binding interfaces rather than a single defined pocket typical of other importin proteins.

### Biochemical insights from our mutations

Through targeted perturbations of the loops, we begin to systematically dissect binding of importins and their cargos from cell-derived samples and learn about the elements regulating importin:cargo interactions. The overlap of our data with others exceeds the field standard for import cargo identification (Supplemental Figure 1E) (14, 41, 42) and along with our HRPF data reassures us that our WT constructs are folding properly (Figure 2C). In all mutant constructs, we observe robust cognate cargo binding by the proteasome, RPS17, Nap1, and TFIIB, which affords strong evidence that our ectopically expressed mutant proteins remain well-folded (17, 36), which is essential for the interpretation of cargo composition changes as being solely due to the focal mutation (Figure 1D, Table 1). The tagged ectopic copies of IPO9 are in a background of the intact endogenous protein and must compete for cargo, further affirming that our mutants are active and reflect real affinity differences to their cargo load. The equivalent protein-level expression of the mutant and WT proteins, with the exception of the ΔTLM mutant, also suggests that the mutant proteins are well-folded, which is critical to the interpretation of competitive redistribution of cargos as a consequence of the mutations.

We enrich for bound proteins from the cytoplasmic pool and largely identify nuclear proteins, emphasizing that these are likely import-competent IPO9-cargo complexes. The steady state distribution of IPO9, greatly favoring the cytosol over the nucleus (Supplemental Figure 6A), also previously noted by microscopy (4, 18) suggests that pore entrance/transport may be rate limiting (67) and IPO9’s nuclear pool rapidly disgorges cargo and exits. This also argues that the cytosol IPO9-bound pool likely reflects its nuclear imported cargo pool. We find that there are diverse modes of binding used by importins; that only a subset of cargos is lost for a given mutation attests to distinct principles by which different cargo classes may be recognized, just as the retention of cargo indicates that a given mutation does not impact that class of protein binding (Figures 4-6). The redistribution of gained cognate cargos upon mutations demonstrates that cargo binding in the cell is the product of competition for limiting IPO9, which is consistent with our HRPF data showing cargo predominantly bound state.

### Interpretations and implications of cognate and secondary cargo binding

Our *in cellulo* mutational scanning approach permits biochemical structure-function dissection using the cell as the test tube. The gain of cognate cargo binding in the mutants could be due to the loss of direct cargo binding for a dominant class of cargos that, through lack of competition, enables other cognate cargo to bind more effectively. The gain of secondary cargos implies two possibilities– loss of repulsive interactions permitting new cargo to be bound that would normally be excluded (gatekeeping is lost), or there is gained affinity due to loss of cognate cargo competition. Only eight proteins found in the WT IPO9 IP (EEF2, RANBP2, AURKB, STIP1, PRPF31, NPM1, P4HB, TSR3) fell below our stringent SAINT threshold but showed affinity to a mutant, indicating that these secondary cargos are largely not present under wild-type conditions and reflect cargos that are normally gatekept from binding to the importin. Loss of the H7 loop specifically results in less stringent recruitment of cargos, as evidenced by decreased enrichment relative to background proteome (Figure 7C). Secondary cargos are very similar in charge/size to cognate IPO9 cargos (Supplemental Figure 6B-C), so discrimination that occurs between cognate and secondary cargos must be on a more subtle scale than solely net charge- or size-based selection. The H7, H8 and H18-19 mutants have a lower percentage of cargo as being annotated as nuclear proteins (Figure 7D), however this is mainly due to ribosomal proteins not being annotated as nuclear, despite known nuclear-localized ribosome biogenesis. The cargo load of IPO9 dynamically changes in each mutant relative to WT as IPO9 is a flexible chaperone protein that accommodates a variety of cargo and uses loop features to discriminate towards and against bound cargo (Figure 7E). When looking at the loop mutants binding profiles, we see that there are overlaps among some secondary cargo additions, most pronounced with the H7 and H8 loop deletions, but also present between the H8 loop deletion and R898E point mutation, indicating that secondary cargos likely have some affinity for the importin but are only able to bind under circumstances where the cognate cargo load is perturbed (Figure 7F). Overall, we can confidently comment on change in abundance 49 of 79 cargos identified in our initial cargo screen (Figure 7G-H). There is an increase in abundance for 18 cargos and a decrease in abundance for 21 cargos with our sets of perturbations.

Intriguingly, there is a large increase in Nap1 without commensurate histone dimer increase in the H8 loop deletion, and there is a loss of histone dimer binding in the R898E and ΔTLM mutants without commensurate loss of Nap1, indicating that the binding of these proteins may be uncoupled at times; this is consistent with previous work noting that Nap1 is able to engage with IPO9 in different orientations, which likely enables its ability to chaperone the histone and promote cargo release (36, 68). While the increase in Ran and RanBP1 binding was unexpected in our H8 loop deletion, we further confirm the important role the H8 loop plays in Ran binding as has been noted for importins generally (6, 9, 11, 49, 50, 62).

### Implications of observed IPO9 properties for nuclear import more broadly

Through our biochemistry-focused mass spectrometry approach, we gain insight into relative cargo affinity for IPO9, mechanistic features of IPO9 loops that may mediate specific cargo recognition, and the complex equilibria of competitive cargo binding that occur upon specific element disruption. Importins are known to have more closed helical structures when traversing the nuclear pore and more open structures, amenable to RanGTP and cargo binding equilibria, in polar environments of cytoplasm and nucleus (69). Navigating the chaotic cytoplasmic milieu to correctly identify cargo is a complex task for the importins. Here, we explore how IPO9-specific loops impact the affinity of the importin towards its various cargos. These loops, present between adjacent HEAT repeats, may allow for further structural flexibility beyond the HEAT repeats flexions to accommodate going through the pore. For cNLS transport, flexible regions adjacent to cargo NLSs impart faster pore trafficking, likely due to increased bound-cargo malleability through opening/closing of the importin (70). Loops on importins may offer a converse mechanism of flexibly tethering cargo to the importin that could make transport more efficient. That we see such a large portion of IPO9 cargos sensitive to loop perturbation reinforces the discriminatory role that loops may play in non-NLS mediated cargo determination (Figure 6H). Inspired by how IDRs effectively sort a variety of proteins in the nucleus, the varied, unstructured loops protruding from the body of importins seem to play an orthogonal, importin-specific role in IPO9 recruitment of cargo.

The vast majority of machine learning-enabled structure predictions of importin:cargo interactions are low confidence ((37), data not shown). Given the dearth of structural information, the high conformational flexibility of importins, and the presumably numerous potential binding modalities of cargo recognition, direct biochemical investigation is critical to begin to understand affinity elements instructing the importin:cargo interactions. We also find that IPO9 engages with sub-groups of cargos (ex. 14-3-3 proteins, PTOV1/MED25, RPS/RPL ribosomal proteins, Transcription Factor ZFs/IDRs, H2A-H2B dimers, including their non-canonical variants, but not other core or linker histones) reaffirm prior work that posits that karyopherins have niches and can co-import fully assembled complexes (14, 42). Interestingly, we do not detect any binding of adaptor importin alpha proteins, sometimes utilized by beta karyopherins for cargo binding, in WT or mutant pulldowns, indicating that IPO9 directly binds its cargo. A few nucleoporin proteins appear in our “Unique to IPO9” WT binding dataset (Supplemental File 1), but they do not rise to the level of our SAINT score threshold, perhaps because they are difficult to immunoprecipitate from cytoplasmic extract due to residence predominantly within the nuclear pore complex. Altered concentration of karyopherins can induce disease (23, 28, 29). We identify cargo binding features that may benefit from increased importin concentration. There is a delicate equilibrium that exists amongst cognate cargo of an importin--affinity range between sub-nanomolar to mid-hundred nanomolar (9, 18, 35), and this, along with a cargo’s relative abundance to other cargos and its importin, ultimately dictates the rate at which a given cargo enters the nucleus after becoming an available cargo. Although we undertook these studies in HEK293 cells, cellular cargo abundance is not likely to be uniform, but rather a feature of any given lineage. While we anticipate that some cargo will be universal, its relative import rate could be a function of competition with lineage specific cargo, such as transcription factors. Another exciting place to look for cargo concentration effects would be during S-phase, when DNA content doubles and histones must be rapidly synthesized and transported into the nucleus – what happens to IPO9’s cargo distribution then? The histones already comprise approximately one fifth of the normal IPO9 cargo load (Figure 1F), but the massive flux of new histone synthesis and import to support DNA replication and chromatin assembly could completely change the cargo load in meaningful and important ways (i.e., relative reduction of actin or a transcription factor import). Conversely, increased cargo expression in disease phenotypes could drastically overtake cargo load and result in decreased import rate of essential cargo, leading to a secondary effect of specific nuclear depletion that would rarely be considered by experimenters. Likewise, overexpression of an importin could induce over-import of cognate cargos or secondary cargos that would otherwise not be imported and result in disease phenotypes through dysregulated compartmental proteome compositions.

### Nuclear release may require more than RanGTP for most IPO9 cargo

Since the discovery of karyopherins in the mid-1990s, the notion that RanGTP induces cargo release has been relatively unchallenged despite a few notable examples of cargo requiring additional factors for release (6, 9, 11, 11, 18, 31, 36, 62, 71, 72). A broader survey has been required to determine which cargos are definitively released with only RanGTP for any importin. In this work, we find that IPO9 cargos are variably sensitive to addition of RanGTP, with most surviving two thirty-minute-long incubations with high concentrations of this general nuclear release factor, arguing that additional factors are likely required to promote the controlled dissociation/deposition of cargos in the nucleus. The affinity for RanGTP among importins is in the low nanomolar affinity regime (<1 - 50 nM)(53, 72) but an importin bound to cargo can have markedly lower affinity for RanGTP (62), arguing that additional factors may be required to induce release of cargo.

The requirement of nuclear release factors is thought to provide a spatio-temporal layer of control over the binding and deposition of cargo. Intra-nuclear concentrations of RanGTP vary relative to proximity to the nuclear pore and chromatin-sensitivity of cargo to RanGTP binding could confer spatial responsiveness to the intra-nuclear environment. IPO9 also carries many cargos that might necessitate more sophisticated nuclear release mechanisms. We observe many proteins that are part of the same complexes: co-transport of such proteins in this form could allow for correct sub-localization of pieces and keep them from mis-assembly with non-cognate partners. Trafficking of proteins to sub-nuclear locations where they act is thought to be essential to drive specific function —this regulation could involve importin-bound factors. In principle, a gradient or spatial sub-localization of a cargo-specific importin release factor could bias this targeting and promote specific release. H2A-H2B histone dimers are a well-established case, requiring a dimer-accepting hexasome or Nap1 to be productively released *in vitro* (18). Somewhat paradoxically, the cNLS containing histone tails sit near the H8 loop of IPO9 and may be required for productive nuclear release *in cellulo* (18, 66, 73). Similarly, one can imagine that releasing the highly basic and non-globular ribosomal proteins without targeting to the nucleolar compartment precisely at the phase of ribosome biogenesis where each becomes specifically embedded in the maturing subunits, could be deleterious to nuclear organization and transcription. The cargos that require the H8 loop are least sensitive to RanGTP binding (Figure 7B) and likely require additional factors to release.

Finally, in considering the possibility for widespread additional cargo release factors, we note that *in vitro* studies implicating RanGTP in cargo release from importins deploy RanGTP at concentrations 2-12 fold above physiological concentrations (18, 32, 65), as do we in this study (10µM). As such, these studies may overestimate the role that individual RanGTP has in dictating nuclear release. Broadly, the identity of additional cargo release factors and how cargos are released in physiological regimes remain central unresolved questions for the field.

## Conclusions

While a simplistic model of cargo specific features that mediate binding to a given importin has been demonstrated for some importin-cargo interactions, such signals are unknown for large subset of importin-cargo interactions. We provide a previously absent *in cellulo* approach to broadly investigate the proteins directly bound to IPO9 and the features that collectively account for binding to more than half of the IPO9 cargos. This framework could be applied to each of the remaining karyopherins, to elucidate their respective cargo cohorts, their potential cargo redundancy, and to define their ortholog-specific cargo recognition principles. Such an endeavor could demystify many aspects of nuclear import, and permit engineering of individual cargo or protein-class-specific modulators of import, as an alternative means of regulating their nuclear function.

## Data availability

All data are contained within the manuscript. The mass spectrometric raw files are accessible at https://massive.ucsd.edu under accession MassIVE MSV000101573 and at www.proteomexchange.org under accession PXD077588. Any inquiries regarding data should be directed to the corresponding author, Alexander Ruthenburg.

## Acknowledgments

We thank Amoldeep Kainth for his technical expertise and advice about mammalian tissue culture and recombinant protein expression and purification.

## Author contributions

A. J. K. and A. J. R. conceptualization; A. J. K., T. C. C., B. M. U. and A. J. R. methodology; A. J. K. and T. C. C. formal analysis; A. J. K., T. C. C., J. K. I. and P. A. S. investigation; A. J. K., resources; A. J. K. and T. C. C. writing; B.M.U. and A. J. R. supervision and editing; A. J. R. funding acquisition.

## Funding and additional information

We gratefully acknowledge support for A. J. K. from the National Science Foundation Graduate Research Fellowship Program and National Institutes of Health T32 training grant (grant no.: T32-GM007197). This study was funded by the National Institutes of Health (grant no.: R35-GM145373) and the Cancer Research Foundation (Fletcher Scholar) grants to A. J. R. The content is solely the responsibility of the authors and does not necessarily represent the official views of the National Institutes of Health.

**Supplemental File 1.** IPO9 cognate cargo identification data

**Supplemental File 2.** HRPF data

**Supplemental File 3.** IPO9 mutant mass spectrometry data

**Supplemental File 4.** RanGTP mass spectrometry data

## Materials and Methods

### Generation of tagged HEK293 FLP-FRT lines

Cells were transfected according to manufacturer of Lipofectamine 2000 (ThermoFisher CAT#11668027) with POG44 recombinase plasmid and appropriate WT or mutant FLP-IPO9 plasmid in 6 well plates. After 5 days, 100ug/mL hygromycin was added to select for successful recombination. Western blotting was performed to confirm presence of tagged full-length recombinant protein.

### Western blotting

HEK293 cells were seeded in 6-well dishes at a density of 1 × 10^6 cells per well and treated with tetracycline (1 μg/mL) for 24 hours prior to harvest. Following induction, cells were washed with ice-cold phosphate-buffered saline (PBS) and lysed in RIPA buffer [50 mM Tris-HCl (pH 7.4), 150 mM NaCl, 1% NP-40, 0.5% sodium deoxycholate, 0.1% SDS, and protease inhibitors] for 30 minutes on ice. Lysates were cleared by centrifugation at 12, 000 × g for 10 minutes at 4°C. Equal amounts of total protein (typically 15–30 μg) were separated by SDS-PAGE on a 10% gel and transferred to PVDF membranes (Millipore) using a semi-dry transfer apparatus according to manufacturer’s instructions. Membranes were blocked for 1 hour at room temperature with 2% w/v blocking agent (Cytiva, RPN2125), then incubated overnight at 4°C with primary antibodies diluted 1:1, 000 in blocking buffer: anti-IPO9 (Abcam, #EPR1352), anti-FLAG (SIGMA, F3165) and anti-GAPDH (Cell Signaling, D16H11) as a loading control. After washing with TBST (Tris-buffered saline, 0.1% Tween-20), membranes were incubated with HRP-conjugated secondary antibody (1:10, 000 dilution in blocking buffer; Sigma, Anti-rabbit IgG A6154 or anti-mouse (ThermoFisher, #31432) for 1 hour at room temperature. Blots were washed extensively and developed by enhanced chemiluminescence (ECL) using Fujifilm LAS4000. Chemiluminescent signals were detected and quantified using the manufacturer’s recommended software.

### Immunoprecipitation-Mass Spectrometry (IP-MS) of FLAG-IPO9 from HEK293 Cells

#### Cell Pellet Growth

Cells were grown as manufacturer recommendation at 37°^C^ with 5% CO_2_ supplemented and 10% Fetal Bovine Serum. 24 hours before collection, 1µg/mL of tetracycline was added to cells to induce expression of the inserted construct. Prior to harvesting, cells were rinsed with 1X PBS and then scraped, spun at 200xg in a Fisher Scientific AccuSpin 1 and flash frozen and stored at −80.

#### Cell Lysis and Fractionation

Whole cell pellets obtained from HEK293 cultures (400 million cells per replicate) were resuspended in ice-cold lysis buffer at a 1:5 (v/v) ratio relative to pellet volume. For quality control purposes, 1% of the lysate was aliquoted for SDS-PAGE/western blotting. Resuspended pellets were mixed gently and incubated at 25°C for 5 minutes. Lysates were centrifuged in a Sorvall Legend XTR at 2, 000 × g for 1 minute at 4°C.). The resulting pellet was resuspended in lysis buffer at a 1:2.5 (v/v) ratio (∼3 mL) and centrifuged again at 2, 000 × g for 1 minute. The pellet was once more resuspended, this time in 1.5 mL of TBP buffer supplemented with 500 μg/mL digitonin and 50 μg/mL lectin for selective permeabilization (39, 79). A final centrifugation was performed at 2, 000 × g for 1 minute, after which the supernatants (cytosolic extracts) were pooled and reserved.

The nuclear pellet was subsequently resuspended in ice-cold TBP-RIPA buffer at a 1:2 (v/v) ratio relative to the original cell pellet (∼2 mL total). Lysis was carried out by end-over-end rotation for 30 minutes at 4°C before centrifugation at 2, 000 × g for 3 minutes at 4°C to extract soluble nuclear proteins. The nuclear extract was further clarified by centrifugation at 4, 700 rpm for 20 minutes at 4°C. Aliquots were taken for immunoprecipitation (IP) input analysis.

For each sample we used 70 μL of 50% bead slurry of anti-FLAG magnetic beads; after equilibrating beads three times with TBP buffer, they were added to the clarified cytoplasmic or nuclear extracts, and the mixture was brought up to a final volume of 15 mL with TBP buffer in a 15 mL conical tube. Immunoprecipitations were performed by rotating the mixture at 4°C for 1 hour. Beads were transferred to 2 mL low retention tubes and washed three times with 1 mL TBP buffer, changing tubes between washes and rotating at 4°C for 5 minutes per wash.

FLAG-tagged proteins were then eluted by adding 300 μL elution buffer (TBP supplemented with 300 μg/mL triple-FLAG peptide, performed in two sequential 150 μL incubations). Each elution was incubated with mixing for 10 minutes at 4°C, and the resulting eluates were combined. Eluted supernatants were collected, and 45 μL aliquots (representing 15% of the IP eluate) were flash-frozen for downstream analysis. For protein quantification and purity assessment, 5% of the sample was loaded on SDS-PAGE gels and visualized with SYPRO Ruby staining, using a BSA standard.

Small scale mutant IPO9 IPs were performed as above but with 40 million cells and proportionally smaller wash/elution volumes.

For RanGTP elutions, 120 μL of 10 μM RanGTP was added and incubated by rotation for 30 minutes at 4°C; this step was repeated for two elutions which were then combined. Beads were then rinsed with 200uL of TBP and then eluted with 3X-FLAG peptide as described above.

Buffer compositions are as follows:

- TBP (20 mM HEPES pH 7.3, 110 mM KOAc, 2 mM MgOAc, 5 mM NaOAc, 0.5 mM EGTA, 2 mM DTT, protease inhibitor cocktail, 8% glycerol) (14)
- Digi/Lectin-TBP Lysis Buffer (TBP supplemented with 500 μg/mL digitonin and 50 μg/mL lectin),
- TBP-RIPA Buffer (TBP base supplemented with detergents per RIPA protocol)
- Elution Buffer (TBP base containing 300 μg/mL triple-FLAG peptide).

### Generation and Analysis of Mass Spectrometry Data

#### Sample Preparation for Binding Partners and Phosphorylation analysis

Samples were reduced by adding 2 µl of 0.2 M dithiothreitol (DTT) and incubated at 57 C for 1 hour. Next, 2 µl of 0.5M iodoacetamide (IAA) was added to the samples to alkylate reduced cysteines. Samples were incubated in the dark at room temperature for 45 mins. Reduced and alkylated samples were loaded onto a NuPage 4-12% Bis-Tris Gel 1.0 mm (Life Technologies). The gel ran for 50 minutes at 200V and stained with GelCode Blue Stain Reagent (Thermo). Gel bands containing proteins were excised for digestion. Excised gel pieces were destained in 1:1 v/v solution of methanol and 100 mM ammonium bicarbonate. The gel pieces were partially dehydrated with acetonitrile and further dried in a SpeedVac concentrator. Dehydrated gel pieces were rehydrated with 100 mM ammonium bicarbonate along with 250 ng of sequence grade modified trypsin (Promega). Digestion proceeded overnight on a shaker at room temperature. The next day, the solution was extracted from the gel pieces and placed into a SpeedVac to dry. The gel was covered with a 1:2 v/v 5% formic acid/acetonitrile extraction buffer and incubated on shaker for 15 mins at 37 C. The extraction buffer was combined with the previously separated solution from the gel and dried in the SpeedVac. Samples were reconstituted with 0.5% acetic acid for offline C18 cleanup. The pH of the reconstituted samples was verified to be <= 2. Samples were loaded onto an equilibrated microspin Harvard apparatus (Millipore) C18 columns. The column was rinsed three times with 0.1% TFA. Peptides were eluted off the column with 40% acetonitrile in 0.5% acetic acid followed by an additional wash of 80% acetonitrile in 0.5% acetic acid. Samples were dried down using SpeedVac and reconstituted in 0.5% acetic acid for LC-MS analysis.

#### Sample Preparation for Hydroxy Radical Protein Footprinting

The hydroxy radical protein footprinting reaction was carried out using three buffers: a standard buffer, a standard-EDTA buffer, and a quench solution. The standard buffer comprised of 25 mM HEPES and 50 mM NaCl at ph 8, while the standard-EDTA buffer additionally contained 330 mM EDTA. The quench solution included 0.1 ng/µl catalase, 510 µM thiourea, and 255 µM methioninamide.

Hydrogen peroxide and Fe(II)-EDTA reagents were prepared fresh at the time of the reaction. A 2.4 M hydrogen peroxide stock solution was prepared in standard buffer. A 1.2 M Fe(II)-EDTA stock solution was prepared by dissolving ammonium iron (II) sulfate hexahydrate in standard-EDTA buffer. Stock solutions were serially diluted in a clear-96 well plate. Concentrations of 1.2 M, 0.60 M, and 0.30 M hydrogen peroxide were generated from the stock solutions, while concentrations of 0.60 M, 0.30 M, and 0.15 M Fe(II)-EDTA solutions were generated from the respective stock solutions. Rows A-D of column 1 contains hydrogen peroxide and rows A-D of columns 2 contain Fe(II)-EDTA at varying concentrations. Row E of columns 1 and 2 contains only standard buffer and standard buffer-EDTA, respectively.

Oxidative footprinting reactions were carried out with Flag-IPO9 and its cargo attached to magnetic beads. To account for the beads themselves quenching the hydroxy radicals, we increased the stock solutions of both hydrogen peroxide and Fe(II)-EDTA, compared to our previous study (38, 48). We created one technical replicate from each biological replicate for a total of 6 replicates (3 biological and 3 technical). 45 µl of the protein-bead mixture was aliquoted to columns A-E and rows 1-5 of a separate dark-96 well plate. The sample plate was mounted to a mixer. 2.5 µl of Fe(II)-EDTA dilutions (column 2 of the dilution plate) were added to the sample plate. Immediately after, 2.5 µl of hydrogen peroxide dilutions (column 1 of the dilution plate) were added to the sample plate. The sample plate was incubated on the mixer for 2 mins at room temperature. The reaction was quenched by adding 50 µl of the quench solution and further mixed for an additional 5 minutes at room temperature.

To remove excess reagent, the samples were mounted on a magnetic rack, and the solutions were extracted and saved. 100 µl of ammonium bicarbonate (pH 8) was added to the beads for reduction, alkylation and digestion. The protein-bead mixtures were reduced using 2 µl of 0.2 M dithiothreitol for 1 h at 57 C, then alkylated using 2 µl 0.5 M iodoacetic acid for 45 mins in the dark at RT. 500 ng of sequencing grade trypsin was added to the samples for overnight digestion on a shaker at room temperature. Digestion was quenched by bringing the pH down to ∼2 with 10% TFA.

#### Mutant Sample Prep For MS/MS analysis

Samples were reduced using 0.2 M dithiothreitol and incubated at 57 C for 1 hour. Next, samples were alkylated using 0.5M iodoacetamide and incubated in the dark at room temperature for 45 minutes. 250 µg of SP3 beads were added to the sample. Proteins were precipitated onto the beads by adding equal volumes of ethanol. The protein-bead mixture was incubated on a shaker at 25 C for 10 mins. The supernatant was removed and saved. The beads were washed with 80% ethanol and incubated on a shaker for 5 mins at 25 C. The supernatant was extracted and saved. The beads were washed in the same manner an additional two times for a total of three washes.

After the last wash, 100 µl of ammonium bicarbonate containing 400 ng of sequencing grade trypsin was added to the samples. Digestion occurred overnight on a shaker at room temperature. Following digestion, the sample was acidified using formic acid, ensuring the ph <= 2. Peptides were loaded on to Evosep tips, preloaded with iRT peptides for MS/MS analysis.

#### Mass Spectrometry methods for Binding Partners and Phosphorylation analysis

Liquid chromatography (LC) separation was performed online with an Easy-nLC 1200 (Thermo Scientific). Peptides were gradient eluted from a PepMap RSLC C18 (ES903, Thermo Scientific) column to an Orbitrap Eclipse Mass Spectrometer using a 1 hour gradient. Solvent A consisted of 2% acetonitrile and 0.5% acetic acid, while solvent B consisted of 80% acetonitrile and 0.5% acetic acid. High resolution MS spectra were acquired in the Orbitrap with a resolution of 120, 000. The AGC target was set to 4e5, with a maximum ion transfer time of 50 ms. The scan range was set to 400 to 1500 m/z. All MS/MS spectra were collected in the Orbitrap with a resolution of 30, 000. The AGC target was adjusted to 2e5 and the maximum ion injection time was set to 200 ms. A 2 m/z isolation widow was used and a normalized collision energy of 32 was used.

#### Mass Spectrometry method for Hydroxy Radical Footprinting

Samples were analyzed using an Evosep One in line with an Orbitrap Eclipse Mass Spectrometer (Thermo Scientific). Five percent of the samples were loaded onto prepared Evosep tips with one injections worth of iRT peptides (indexed retention time peptides, Biognosys). Evosep tips were rinsed a total of two times with 0.1% formic acid before loading onto the Evosep for analysis. The tips remained wet until they were analyzed. Peptides from each sample were eluted using the 15SPD (88 min gradient) Evosep method. Solvent A consisted of 0.1% formic acid, while solvent B consisted of 0.1% formic acid in acetonitrile. Mass spectrometer settings are the same as the settings for binding partners and phosphorylation analysis. However, the dynamic exclusion was reduced from 30 seconds to 5 seconds to enhance sampling of oxidized isoforms.

#### Data Analysis for Binding Partners

To evaluate the binding partners for IPO9, the MS/MS spectra was searched using Sequest within Proteome Discoverer 1.4. The precursor mass tolerance was set to 10 ppm while the fragment mass tolerance was set to 0.02 Da. A maximum of 2 missed cleavages were allowed. A 1% FDR cutoff was used for peptide spectra assignment. Oxidation on methionine (M), deamidation on asparagine (N) and glutamine (Q), and phosphorylation on serine (S), threonine (T), and tyrosine (Y) were included as variable modifications. Carbamidomethyl on cystines (C) were included as fixed modification. The data was further filtered by only considering peptide with High confidence. A minimum of 0.15 Dela Cn was used to further filter peptide-spectra matches. Single hit proteins were filtered out by requiring at least 2 peptides per protein. Significant binding partners were identified using SAINT method. A SAINT score cutoff of 0.9 was used to confidently assign binding partners (Supplemental Files 1, 3 and 4).

#### Data Analysis for Phosphorylation Site identification

MS/MS spectra were searched using Byonic implemented in PMI-Byos (Protein Metrics). The precursor and fragment mass tolerance was set to 10 ppm. A maximum of 2 missed cleavages were allowed. Carbamidomethyl on cystines was set as a fixed modification. Oxidation on methionine/tryptophan were set to variable rare 1 modification. Deamidation on asparagine/glutamine was set to variable rare 1. Phosphorylation on serine\threonine\ tyrosine were set to variable common 2. GlyGly modifications on lysines were set to variable common 1. All phosphorylated spectra were manually inspected. A byonic score of 300 was used as a cutoff for phosphorylated peptide-spectrum matches.

#### Data Analysis for Hydroxy Radical Footprinting

MS/MS spectra were searched using the oxidative footprinting node of Byos (Protein Metrics) against a Uniprot Homo sapien database. Spectra were searched fully tryptic allowing for two missed cleavages, maximum precursor mass of 10000 Da, 10 ppm precursor tolerance, 20 ppm fragment ion tolerance, 1% FDR, manual score cut of 300, two common modification maximum, and one rare modification maximum. Trioxidation at cysteine, dioxidation at cysteine/methionine/tryptophan, and oxidation at isoleucine/lysine/leucine/glutamine/valine were set as common 1 modification. Oxidation at methionine/cysteine/aspartic acid/phenylalanine/histidine/phenylalanine/tryptophan/tyrosine/proline was set as a common 2 modification. pyroGlu at N-term glutamine and N-term glutamic acid was set as a rare 1 modification. All peptides were manually validated and integrated using the Byos platform. Oxidation percentages were determined by subtracting each dose by the oxidation percentages from the samples where no oxidizing reagents were added (controls). Each dose represents the average and standard deviation of five replicates.

#### Data Curation and Visualization

GraphPad Prism was used for data visualization (80). BioVenn was used for venn diagrams (81). ChimeraX was used to visualize the IPO9 protein (82). Deeploc2.1 was used to categorize IPO9 cargo(45). AIUPRED was used for importin disorder scores (76). Consurf-DB was used to analyze IPO9 conservation amongst homologs (77). Post-translational modification data found on Phosphosite plus (61). Cargo molecular weight and isoelectric point data were calculated using the Expasy compute MW/PI tool (78).

#### Cloning, Expression and Purification of Proteins

His-IPO9-GST was cloned from two G-blocks into a pGEX-6P plasmid. Mutant IPO9 constructs were cloned via Gibson assembly reactions with the parent plasmid. Protein expression constructs were transformed into Rosetta 2 (DE3) pLysS competent cells and grown in 1L LB cultures containing 25 µg/mL chloramphenicol and 100 µg/mL carbenicillin at 37°C until an optical density at 600 nM (OD600nm) measured ∼ 0.6. Upon reaching this OD600nm, cultures were cooled to 18°C then induced with 1mL of .4 mM isopropyl-β-D thiogalactopyranoside (IPTG) then shaken at 18°C for 18 hours. Cultures were then harvested by centrifugation in a Thermo Sorvall Lynx 6000 at 4000g for 20 min at 4°C. Cell lysis via 3 passes through the Avestin EmulsiFlex-C3 at 15, 000 PSI then clarified by centrifugation on Sorvall Lynx 6000 at 30, 000g for 20 min at 4°C. Lysate was filtered through 0.45 µM before being loaded onto FPLC via 50mL Superloop. IPO9 was purified to homogeneity by first a 5mL HisTrap HP Nickel column (Cytiva, #17524801) followed by GSTrap 4B 5mL column (Cytiva, #28401747). Subtractive nickel chromatography was used to concentrate the protein and change the protein into TBP buffer and then dialyzed into TBP to remove imidazole from buffer.

For the recombinant HRPF procedure, the GST tag was cleaved with R3C overnight post-GSTrap column. A S200 sizing column was then ran, followed by overnight dialysis into 20mM HEPES pH 7.6, 150mM NaCl, 8% glycerol.

RanQ69L sequence was ordered via GBlock and cloned into background Pet16B plasmid containing a 10X-His and CL7 tag. Plasmid expression was induced at .6OD with .4mM IPTG in Rosetta 2 (DE3) pLysS cells for 4 hours at 37 degrees Celsius. Cells were spun at 4000xg on Themo Sorvall Lynx 6000 for 20 minutes to pellet and then flash frozen and stored at −80 degrees Celsius freezer until use. Lysis of pellet in lysis buffer (20mM Tris pH=7.5, 100mM NaCl, 10% glycerol, .5mM PMSF, 1uL benzonase, 1X PI cocktail) and lysed vis French press. Lysate was clarified via centrifugation 5000*g for 20 minutes in Sorvall Lynx 6000 rotor. Lysate was then filtered through a .45µM filter then loaded onto FPLC via 50mL Superloop. Protein was purified via nickel affinity chromatography followed by anion exchange and tag cleavage via R3C. Ran was loaded with GTP as described previously (35, 83, 84) (GTP was added for 2 hours on ice at 4 degrees with 5mM MgCl2 in the reaction, then MgCl2 was increased to 20mM with 10mM EDTA, then flash frozen for later use.)

Cargo proteins were acquired from Addgene and induced at 0.6 OD then expressed for 4 hours at 37 degrees 225 rpm, followed by pelleting and lysis as described above. Cargo were lysed via Emulsiflex and purified on nickel resin (Qiagen Ni-NTA) and eluted in 1mL 1M imidazole containing HBS.

pNH-TrxT-G-S18 was a gift from Ivanhoe Leung (Addgene plasmid # 127827; http://n2t.net/addgene:127827; RRID:Addgene_127827)(85)

CPSF5A was a gift from Nicola Burgess-Brown (Addgene plasmid # 42353; http://n2t.net/addgene:42353; RRID:Addgene_42353)

H2A-H2B purification and dimer refolding were performed as described previously (85).

#### Recombinant Cargo pulldowns

200uL of nickel elution of each cargo were used per pulldown. 5uL of magnetic GST beads were used per reaction. GST beads were equilibrated in binding buffer + reducing agent, then IPO9-GST (WT or mutant) was added for 30 minutes at room temperature, rotating. After 30 minutes, 3×1mL washes of the beads to rinse away unbound bait. Following this, cargo was added for a total reaction volume of 400uL. Reactions rotated for 45 minutes at room temperature and then were followed by 1mL washes with 5 minutes rotation. Beads were resuspended in 50uL of BB and 10uL of 6X Loading dye, then boiled to elute. 10 or 15% SDS-PAGE gels were loaded with 15uL of elution.

#### Fluorescence Polarization

Recombinant IPO9 was serially diluted in triplicate from 3000nM to 5nM in a 20mM Tris pH=8.0, 150mM NaCl, 10% glycerol containing solution with 20nM WT and mutIFNE ssRNA labeled with 5’ FITC and measured at excitation/emission wavelengths of 485 nm/535 nm on a TECAN Infinite 200 Pro.

---

**Supplemental Figure 1.**
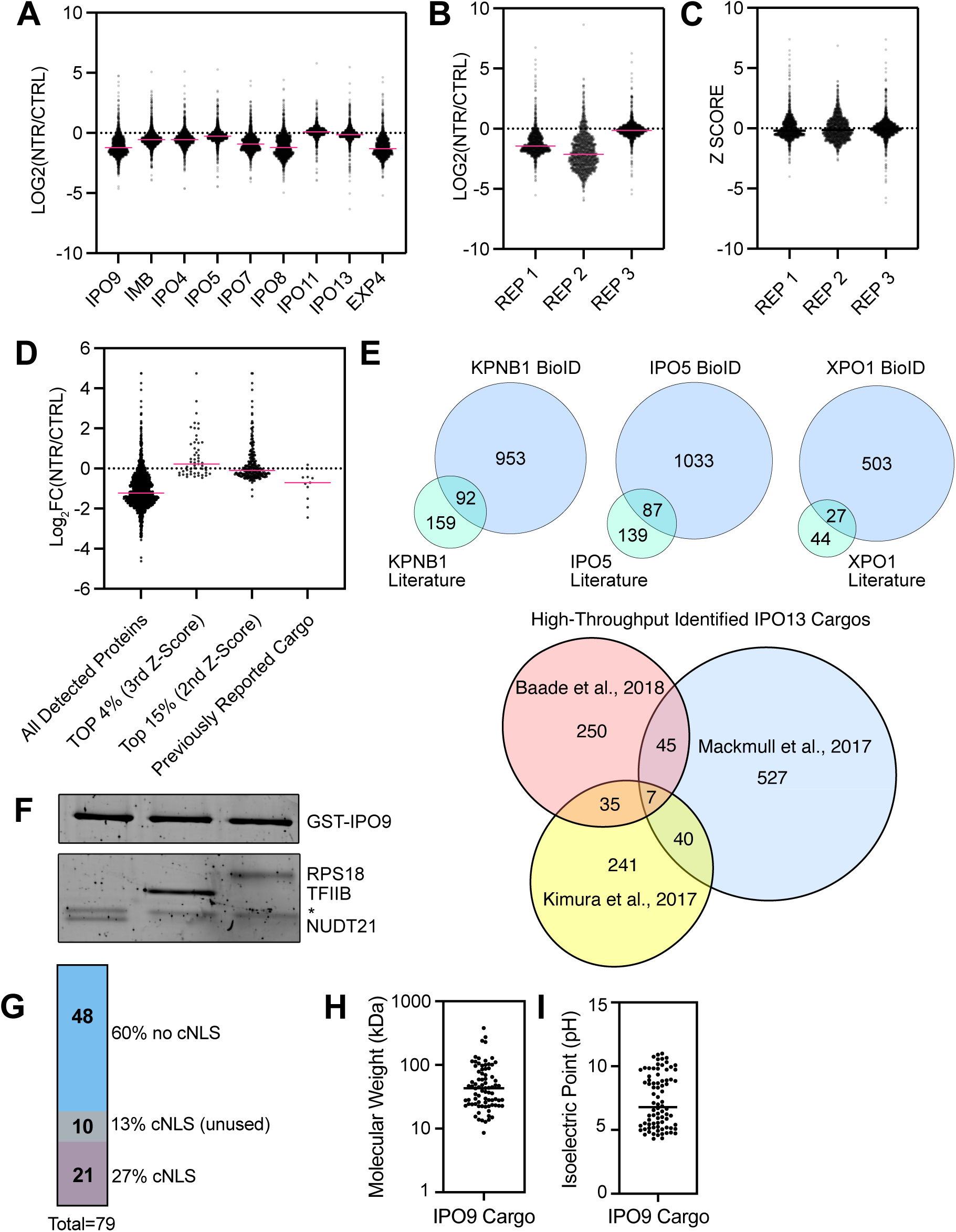
Comparison to previous importin•cargo interaction data and further investigation into identified IPO9 cargo characteristics. (A) Graph detailing the Kimura 2017 group’s log2 fold-change in all cargos measured for each importin. NTR means nuclear transport receptor, aka karyopherins. Note variation from baseline (0 fold change) for IPO9, IPO7, IPO8, and EXP4/XPO4. This variation may be expected for XPO4 as it is a biportin, so one would expect less nuclear signals of some cargoes. (B) Log2 fold-change for IPO9 experiment by Kimura 2017 group broken out by replicate. Large variation between replicates. (C) Transformation of Log2 fold-change values into Z-scores masks the variation seen in the raw data. (D) When looking of the Log2 Fold changes of cargo identified as significant (top 4% 3^rd^ z-score n=53, top 15% 2^nd^ z-score n=254), one can see that many of these cargos are not enriched in the +IPO9 samples. Import scores of previously reported cargo are similarly variable. (E) Recapitulation of Venn diagrams from Mackmull 2018 paper showing overlap of identified cargo with existing dataset. (F) SYPRO Ruby stained 10% SDS-PAGE gel depicting putative cargo pulldowns with highly purified recombinant factors: GST-IPO9 and NUDT21, TFIIB, or RPS18 (*corresponds to nonspecific protein present in GST-IPO9 protein preparation). (G) Plot depicting the fraction of cargo that have mono- or bi-partite NLSs present in their amino acid sequence by NLSstradamus (46). (H) Chart depicting molecular weight of identified cargos (78). (I) Chart depicting isoelectric point of identified cargo. IPO9, importin 9.

**Supplemental Figure 2.**
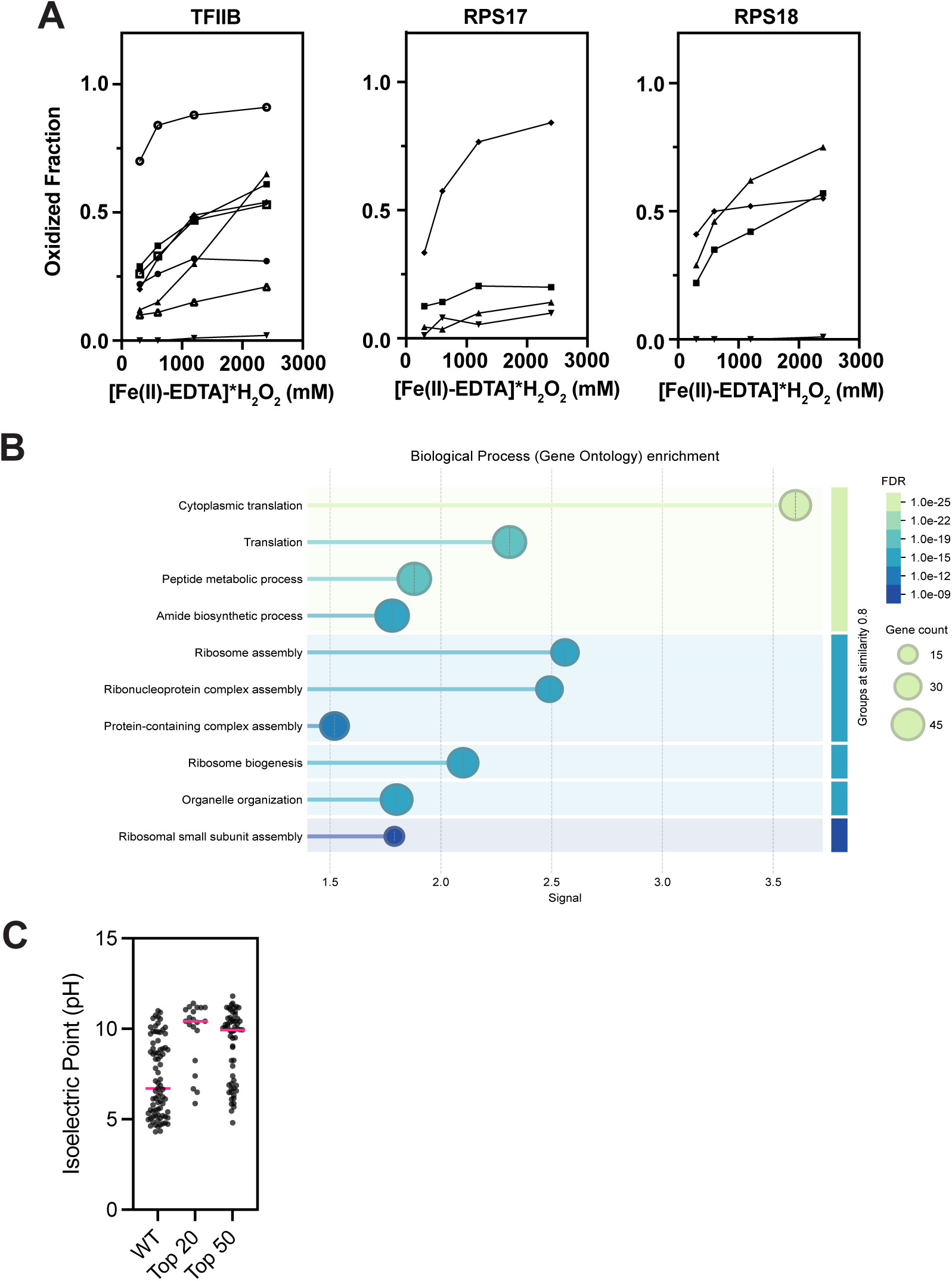
Ox-FT identified peptides of cargo proteins. (A) Peptide oxidation fraction for three different cargo proteins.

**Supplemental Figure 3.**
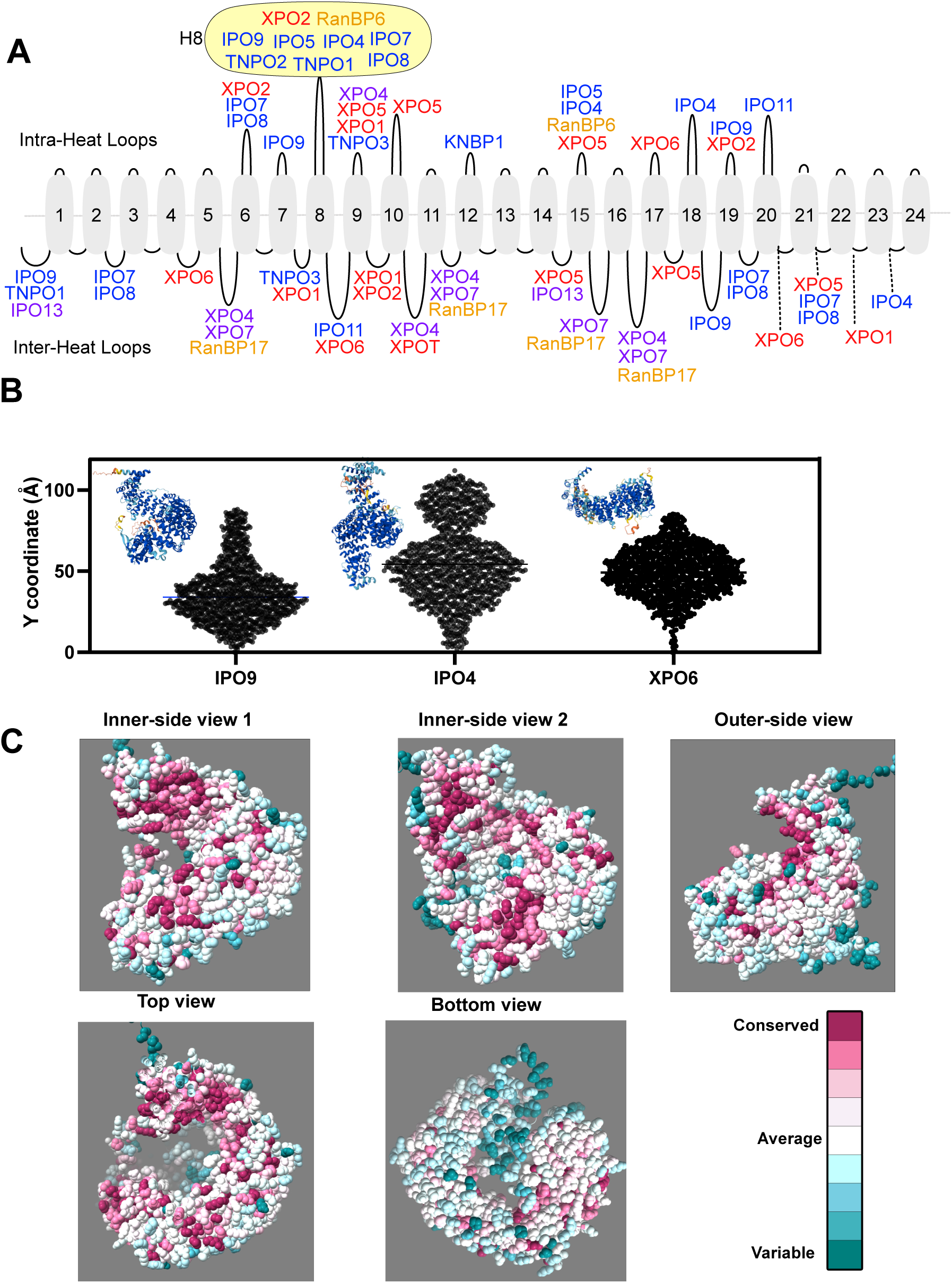
Radius and pitch of importins is variable despite HEAT repeat conservation. (A) Depictions of loops of all karyopherins spread along an example karyopherin with 24 heat repeats. Importins are blue, exportins red, biportins purple, unknown directionality orange. Length of loop does not correspond to real loop length. H8 loop having proteins are highlighted in yellow. (B) Alphafold predicted structures of IPO9, IPO4 and XPO6 overlayed on graph depicting Y-axis (height) coordinate of each importin mapped for each residue. Note how IPO4 is taller than either IPO9 or XPO6. (C) CONSURF scores mapped onto PDB:6N1Z (18), shown at different angles.

**Supplemental Figure 4.**
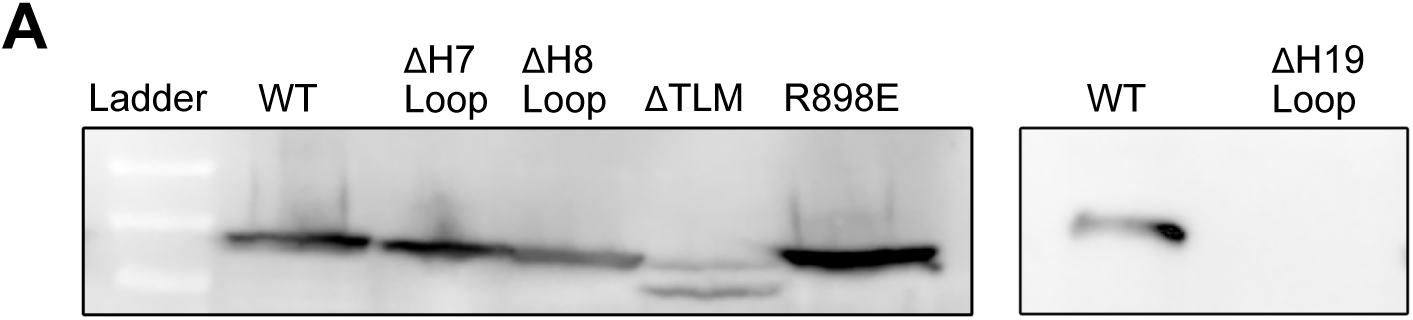
Confirmation and quality control of mutant IPO9 FLP-FRT lines. (A) Western blot using anti-FLAG antibody to probe for expression of mutant FLAG-tagged IPO9 lines (WT, ΔH7, ΔH8, ΔTLM, R898E). Note that dH19 line did not noticeably express.

**Supplemental Figure 5.**
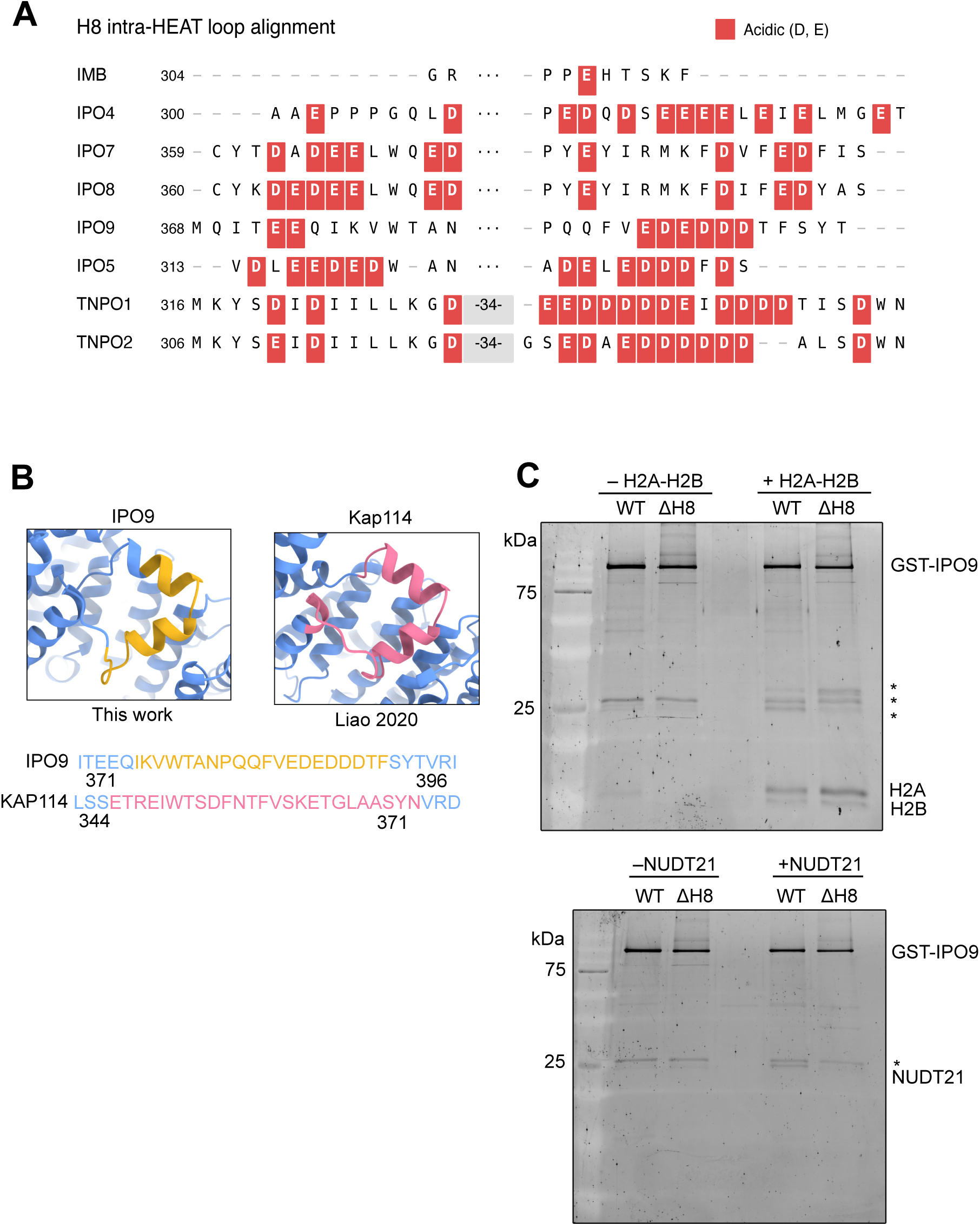
Understanding cargo release features of IPO9. (A) Conservation analysis of the H8 loops of importins aligned with MUSCLE (B) Visual and sequence comparison of mutants made by our group (yellow, no linker added), the Chook lab (yellow and red, replaced with SGSTGGSGS), and the Hsieh group with Kap114 (pink, no linker added). Perturbations mapped onto Alphafold predicted structures for IPO9 and Kap114. (C) Full gel of recombinant IPO9 pulldowns for H2A-H2B and NUDT21.

**Supplemental Figure 6.**
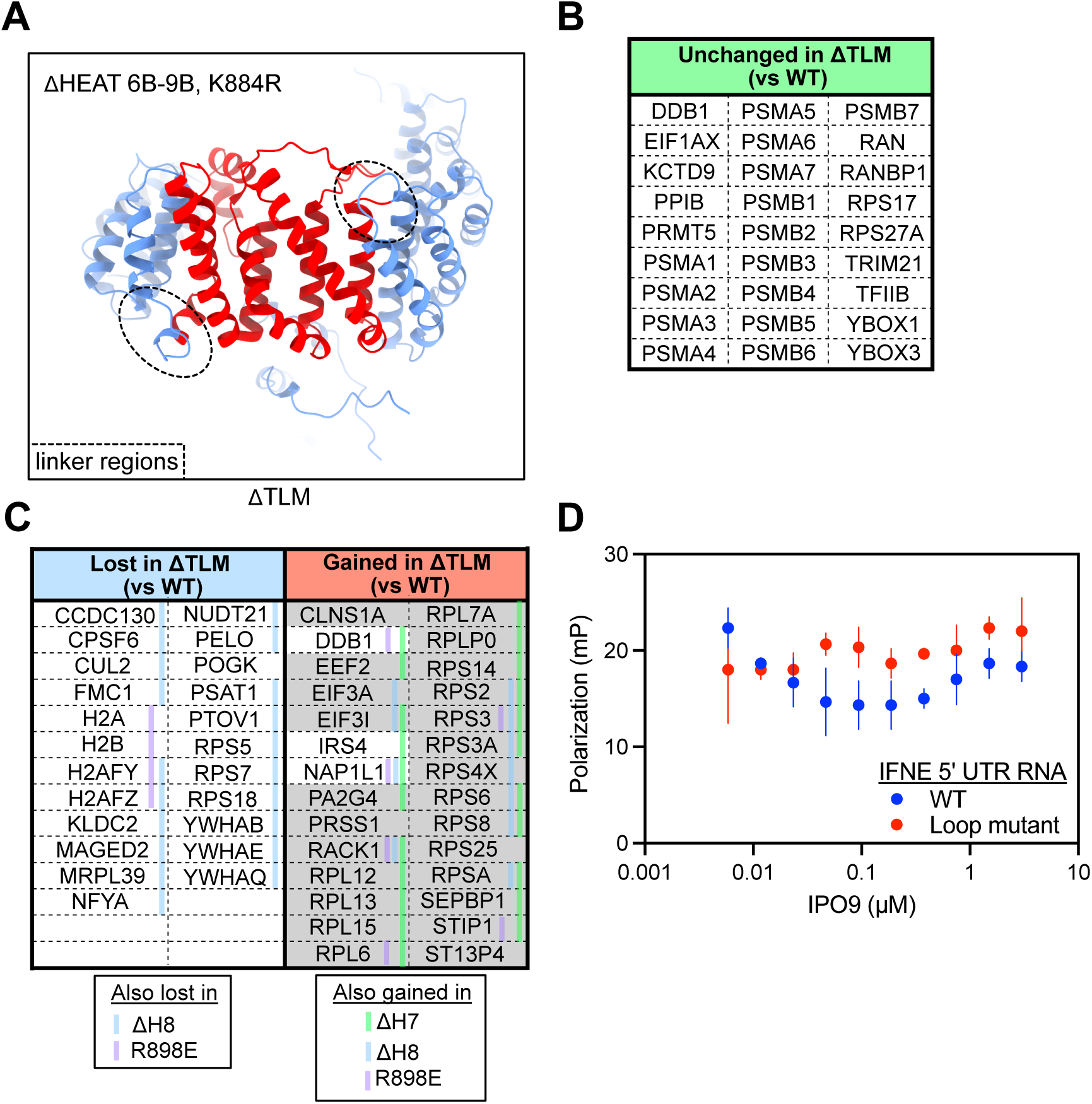
ΔTLM confirms individual loop perturbations; IPO9 does not bind RNA. (A) Structural view highlighting unstructured regions present in the ΔTLM structure that allow for proper folding of regions adjacent to the middle section deletion. (B) Cognate cargos unchanged in ΔTLM mutant (C) Cognate cargos lost and ΔTLM cognate and secondary (grey) cargos gained in the ΔTLM mutant, with annotations of other mutants cargo noted in key. (D) Investigation into IPO9 RNA binding activity. 5’FAM labeled WT or mutant IFNE 5’ UTR RNA sequence (25) (20nM) was assayed over a (0-3000nM) range of recombinant IPO9 in a fluorescence polarization assay (n=3).

**Supplemental Figure 7.**
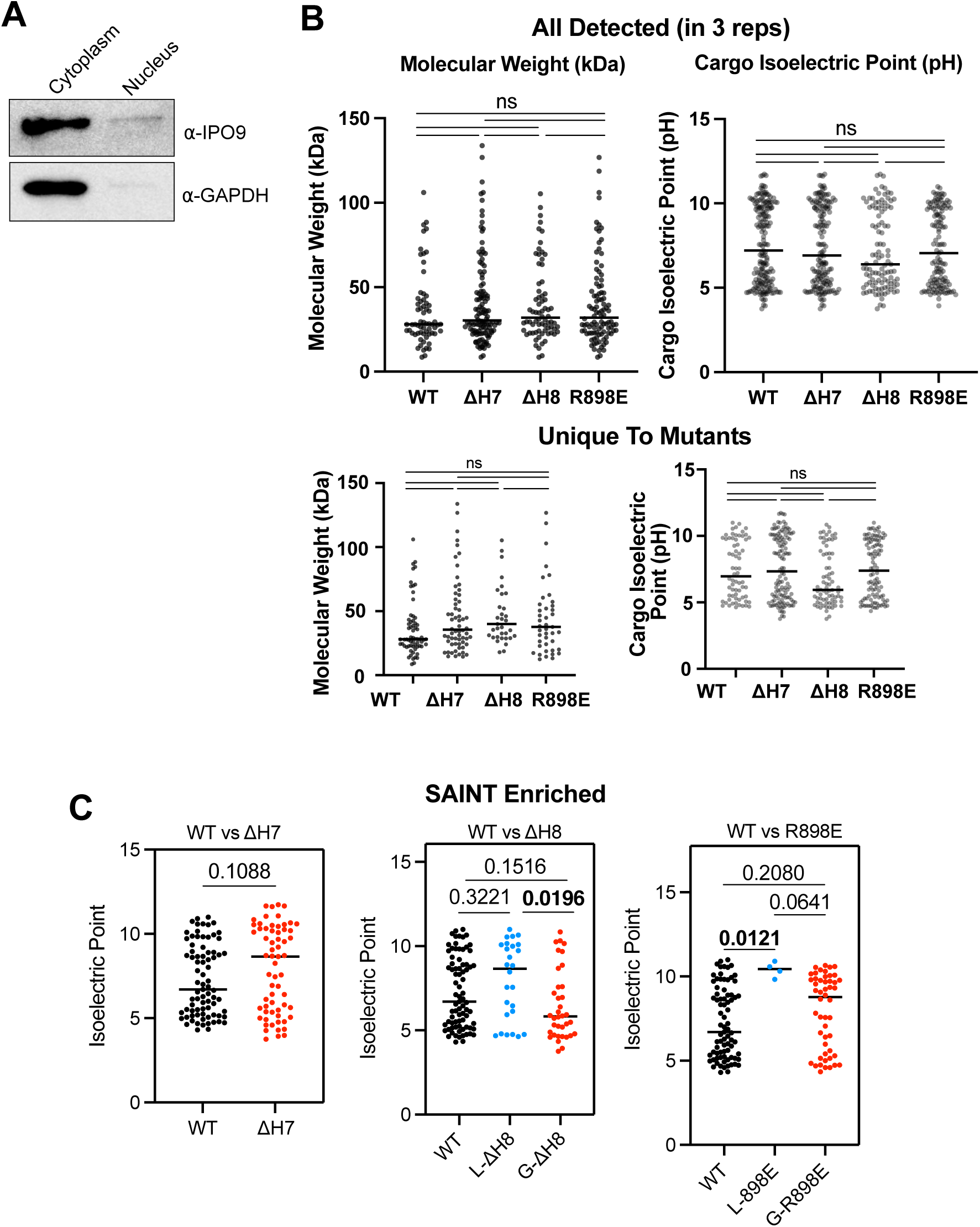
IPO9 localization and comparison of cargos bound by mutants. (A) Western blot of IPO9 from nucleus and cytoplasm fractionated HEK293 cells. GAPDH as fractionation control. (B) Cargo isoelectric point and molecular weight comparisons of all cargo detected in a given ectopic expression pulldown. One dot equals one cargo. Plotted for total cargos seen in each pulldown and for cargos that are unique to the mutant relative to the WT. (C) Isoelectric points of WT cargo versus cargos that are SAINT enriched >0.9 in each mutant pulldown.

**Supplemental Table 1.** PANTHER GO term analysis for cargos enriched SAINT >0.9.

| GO Term | GO ID | Ref | Hits | Fold Enrichment | FDR |
| --- | --- | --- | --- | --- | --- |
| *Proteasome & Protein Degradation* |  |  |  |  |  |
| Proteasome-mediated ubiquitin-dependent protein catabolism | GO:0043161 | 244 | 13 | 14.06 | 2.55E-08 |
| Modification-dependent protein catabolic process | GO:0019941 | 379 | 14 | 9.75 | 1.64E-07 |
| Ubiquitin-dependent protein catabolic process | GO:0006511 | 370 | 13 | 9.27 | 8.79E-07 |
| Proteolysis involved in protein catabolic process | GO:0051603 | 452 | 15 | 8.76 | 1.80E-07 |
| Protein catabolic process | GO:0030163 | 467 | 15 | 8.47 | 1.77E-07 |
| Proteasomal protein catabolic process | GO:0010498 | 299 | 13 | 11.47 | 1.62E-07 |
| *Chromatin & DNA* |  |  |  |  |  |
| Heterochromatin formation | GO:0031507 | 76 | 4 | 13.89 | 4.03E-02 |
| DNA-templated transcription | GO:0006351 | 140 | 5 | 9.42 | 4.01E-02 |
| Chromatin organization | GO:0006325 | 266 | 9 | 8.93 | 2.67E-04 |
| Chromatin remodeling | GO:0006338 | 222 | 7 | 8.32 | 5.65E-03 |
| *RNA Processing & Metabolism* |  |  |  |  |  |
| mRNA alternative polyadenylation | GO:0110104 | 2 | 2 | >100 | 3.91E-03 |
| rRNA metabolic process | GO:0016072 | 147 | 5 | 8.97 | 4.74E-02 |
| mRNA processing | GO:0006397 | 181 | 6 | 8.75 | 1.54E-02 |
| mRNA metabolic process | GO:0016071 | 255 | 7 | 7.24 | 1.27E-02 |
| RNA biosynthetic process | GO:0032774 | 619 | 14 | 5.97 | 3.66E-05 |
| RNA metabolic process | GO:0016070 | 742 | 15 | 5.33 | 4.76E-05 |
| RNA processing | GO:0006396 | 479 | 9 | 4.96 | 1.76E-02 |
| *Broader Metabolic Processes* |  |  |  |  |  |
| Nucleic acid biosynthetic process | GO:0141187 | 639 | 14 | 5.78 | 4.73E-05 |
| Nucleobase-containing compound biosynthetic process | GO:0034654 | 764 | 14 | 4.83 | 3.11E-04 |
| Nucleic acid metabolic process | GO:0090304 | 1060 | 18 | 4.48 | 3.37E-05 |
| Macromolecule catabolic process | GO:0009057 | 581 | 16 | 7.27 | 3.46E-07 |
| Catabolic process | GO:0009056 | 1007 | 16 | 4.19 | 3.25E-04 |
| Gene expression | GO:0010467 | 1159 | 17 | 3.87 | 3.84E-04 |
| Protein metabolic process | GO:0019538 | 1409 | 20 | 3.75 | 7.76E-05 |
| Macromolecule metabolic process | GO:0043170 | 2499 | 37 | 3.91 | 1.10E-10 |
| Nucleobase-containing compound metabolic process | GO:0006139 | 1318 | 18 | 3.60 | 4.69E-04 |

Ref = number of genes in reference list; Hits = number of genes in uploaded list FDR = false discovery rate corrected p-value (Fisher's exact test)

